# An allosteric pocket in K_V_1.3 defines a distinct chemical space for immunomodulator design

**DOI:** 10.64898/2026.07.06.736759

**Authors:** Guoqing Luo, Xin Zhang, Huixin Xia, Ziyue Wei, Zheyang Zhang, Jie Sun, Zixuan Zhang, Yi Peng, Han Liu, Xuemeng Huang, Peng Cao, Mingqiang Rong, Ye Yu, Cheng Tang

**Affiliations:** The National and Local Joint Engineering Laboratory of Animal Peptide Drug Development, College of Life Sciences, Hunan Normal University, Changsha 410081, China; Peptide and small molecule drug R&D platform, Furong Laboratory, Changsha 410078, China; Key Laboratory of Preclinical Study for New Drugs of Gansu Province, School of Basic Medical Sciences, Lanzhou University, Lanzhou, China; Hospital of Integrated Traditional Chinese and Western Medicine, Nanjing University of Chinese Medicine, Nanjing 210023

## Abstract

K_V_1.3 is a validated autoimmune drug target, yet all disclosed inhibitors converge on a narrow set of conserved binding sites, suffering from poor selectivity or clinical failure. Whether K_V_1.3 harbors a distinctive, druggable pocket amenable to selective targeting has remained unknown. Here we report MPiN, a highly selective, state-dependent K_V_1.3 inhibitor with in vivo efficacy in a mouse psoriasis model, which engages a previously uncharacterized extracellular allosteric pocket framed by the PP1-PP2 turret loops, the pore helix and the outer S5/S6 helices, acting through a bidirectional pore-to-sensor coupling that simultaneously constricts the selectivity filter and facilitates the voltage sensor toward activation. Unexpectedly, despite the extensive structural conservation of this pocket across K_V_1 paralogs, subtype selectivity is dictated by the peripheral residues G427 and H451, which define pocket geometry without directly contacting the ligand, thereby establishing a geometry-driven “non-contact selectivity” mechanism. By opening an unrecognized, structurally distinct chemical space on K_V_1.3 and redefining how selectivity is achieved within a conserved channel family, this work lays the structural and conceptual foundation for rational, structure-guided design of next-generation K_V_1.3 immunomodulators.

## Introduction

The voltage-gated potassium channel K_V_1.3 is a homotetramer of four α-subunits, each spanning six transmembrane helices (S1–S6)^1^. Depolarization drives the S4 helix outward through a gating pore formed by S1–S3; this movement allosterically opens the pore domain (S5–S6 and its linker), enabling selective KL conduction^2–4^. Sustained depolarization instead triggers C-type inactivation through rearrangement of the selectivity filter ^5,6^. As a gated KL transporter, K_V_1.3 is central to immune regulation and tumor biology ^7,8^. It is selectively enriched in effector memory T (T_EM_) cells and largely absent from central memory T (T_CM_) cells^9–11^. KL efflux through K_V_1.3 sustains Ca²L entry via calcium-release activated calcium (CRAC) channels, driving Ca²L/NFAT (nuclear factor of activated T cells)-dependent T_EM_ activation - a core pathogenic axis in autoimmune diseases (ADs) ^10^. In mitochondria, K_V_1.3 is inhibited by the pro-apoptotic protein Bax, triggering membrane depolarization, cytochrome c release, and apoptosis ^12^. Many cancers subvert this axis: K_V_1.3 is upregulated, Bax is downregulated, and tumor cells evade death^8,13^. K_V_1.3 has therefore emerged as a premier therapeutic target - selectively silencing pathogenic T_EM_ responses in ADs while sparing normal immunity, and offering a compelling entry point for cancer therapy ^10,11^.

Two pharmacological categories of K_V_1.3 inhibitors have been developed, each with distinct mechanisms and therapeutic profiles. Peptidic inhibitors, largely derived from animal venoms, act either by trapping the S4 helix to impede gating - exemplified by the spider toxin MrVIII ^14^ - or by occluding the K^+^ pore through binding to the outer vestibule, including ADWX-1^15^, ShK^16^, AcK1^17^, BmK1^18^, ChTX^19^, MgTX^20^, and OSK1^21^. Dalazatide (ShK-186), engineered from the sea anemone toxin ShK for enhanced K_V_1.3 selectivity and stability^22^, advanced into clinical trials but was subsequently suspended despite its success in the Phase 1b stage. By contrast, the peptide-based K_V_1.3 inhibitor SI-544 continues in active clinical development, supported by positive Phase 1b results (NCT05383378;NCT06191042). Poor oral bioavailability remains the principal limitation of this class. Small molecules were developed to overcome these constraints, offering improved pharmacokinetics, lower production cost, and greater chemical stability. Reported compounds include synthetic scaffolds (UK-78282, WP1066, WIN17317-3, CP-339818)^23–26^, repurposed drugs (clofazimine and verapamil)^27,28^, and natural products (correolide, psoralen, loureirin B)^29–31^. In contrast to peptidic outer-pore blockers, these molecules all engage the inner cavity of K_V_1.3 - a region highly conserved across the voltage-gated K^+^ channel superfamily^32^ - and consequently exhibit limited subtype selectivity, an obstacle that has stalled their clinical development. A notable exception is Psora-4, which engages a unique transmembrane side pocket besides the central cavity ^33^. Further optimization of Psora-4 yielded PAP-1, another highly selective K_V_1.3 inhibitor with preclinical therapeutic potential across multiple diseases^34–37^. However, despite completing a Phase 1b trial, its clinical translation remains uncertain (NCT01743118). DES7114, a K_V_1.3-specific inhibitor designed de novo via all-atom conventional molecular dynamics (CMD) simulations to trap the channel in its inactivated state, has advanced to Phase II trials - the highest global R&D status among K_V_1.3-targeting drugs ^38^(NCT06176768). Collectively, among documented K_V_1.3 inhibitors, DES7114 exhibits high druggability and operates through an undisclosed mechanism, whereas the remainder converge on a narrow set of conserved binding sites, suffering from poor selectivity or outright clinical failure. Identifying selective K_V_1.3 inhibitors that engage previously unrecognized binding pockets and act through distinct mechanisms will pave the way for next-generation K_V_1.3 therapeutics.

Through subtractive screening of a structurally diverse compound library, we identify MPiN, a highly selective small-molecule K_V_1.3 inhibitor with pronounced therapeutic efficacy in a murine autoimmune disease model. MPiN engages a previously uncharacterized extracellular pocket that is conserved across the K_V_ channel superfamily, and inhibits K_V_1.3 in a state- and allosteric-dependent manner. Selectivity for K_V_1.3 is dictated by two channel-specific residues, which shape pocket geometry without directly contacting the ligand, a selectivity mechanism distinct from all reported K_V_1.3 inhibitors. These findings established an allosteric paradigm of selective K_V_1.3 antagonism and revealed a distinct druggable pocket for the design of next-generation immunomodulators.

## Results

### Subtractive screening identified a potent and highly-selective K_V_1.3 inhibitor

The high degree of sequence conservation across the voltage-gated potassium (K_V_) channel superfamily has made the development of subtype-selective inhibitors particularly challenging, as most reported inhibitors target conserved binding hotspots and consequently exhibit extensive cross-reactivity. To overcome this limitation, we devised a rational subtractive screening strategy to identify K_V_1.3-selective inhibitors that engage structural features unique to K_V_1.3 rather than the conserved sites commonly exploited by known K_V_ channel inhibitors. A structurally diverse library of 4,208 compounds was first screened against K_V_1.3 channels, and compounds producing >20% inhibition at 10 μM were subsequently counter-screened against the closely related K_V_1.5 channel (**Fig.1A**).

**Figure 1.**
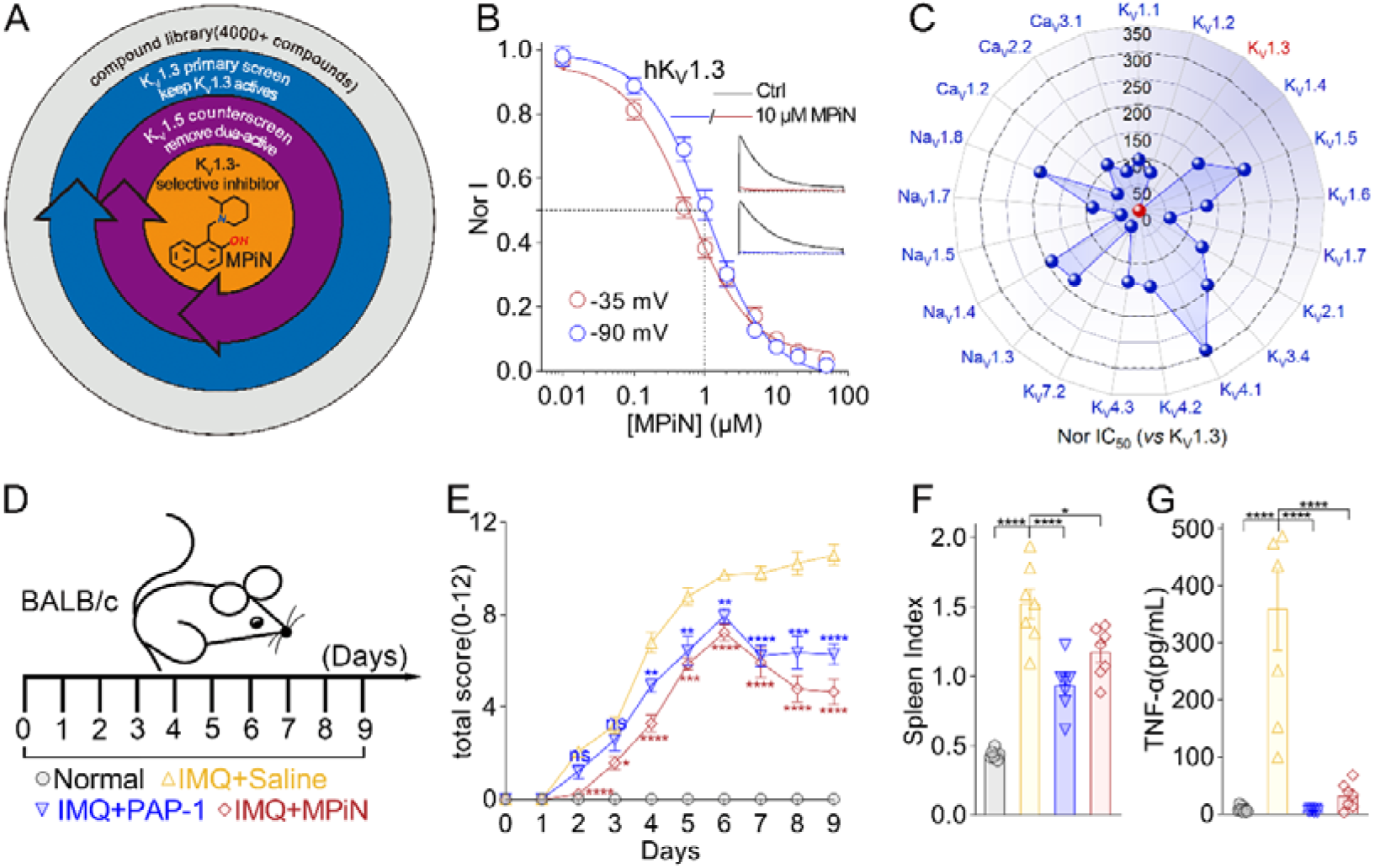
Electrophysiological characterization of MPiN as a novel K_V_1.3-selective inhibitor. (A) Subtractive screening strategy: library compounds were first tested against K_V_1.3, and hits were counter-screened on K_V_1.5 to eliminate dual-active compounds. MPiN emerged as the sole K_V_1.3-selective hit. (B) Representative traces (inset) and dose-response curves of MPiN inhibiting K_V_1.3 currents, yielding IC_50_ values of 0.6 ± 0.1 μM at −35 mV and 1.1 ± 0.1μM at −90 mV (n = 7-12). (C) IC_50_ values of MPiN against off-target ion channels normalized to K_V_1.3, demonstrating high selectivity for K_V_1.3 (n =5-9). (D) Experimental design showing induction of psoriasis by imiquimod (IMQ) in BALB/c mice and the treatment regimen. (E) Total Psoriasis Area and Severity Index (PASI) score for animals in each group along time (n =7). (F)-(G) Spleen index (F) and serum TNF-α (G) level at D9 (n =7).

This screening campaign identified 20 chemically distinct K_V_1.3 inhibitors that inhibited channel currents by 26-89% (**Supplementary Fig.1A,B**). Strikingly, only one compound, designated MPiN, satisfied our stringent selectivity criteria, exhibiting both high potency and a pronounced preference for K_V_1.3 over K_V_1.5 (**Fig.1A and Supplementary Fig.1A-C**). Electrophysiological characterization revealed that MPiN inhibited K_V_1.3 currents with IC_50_ values of 1.1 ± 0.1 μM and 0.6 ± 0.1 μM when channels were held in predominantly closed (−90 mV) and inactivated (−35 mV) states, respectively (**Fig.1B**). Inhibition developed rapidly upon compound application and recovered readily following washout, consistent with reversible binding (**Supplementary Fig.1D**).

Comprehensive profiling across multiple voltage-gated potassium, sodium, and calcium channels demonstrated remarkable selectivity for K_V_1.3, with MPiN exhibiting ∼32-320-fold lower potency toward all tested off-target channels (**Fig.1C and Supplementary Fig.2**). Importantly, MPiN also showed robust efficacy in an imiquimod-induced mouse psoriasis model, significantly attenuating disease progression, reducing splenomegaly, and suppressing plasma TNF-α levels, with efficacy comparable to that of PAP-1, a K_V_1.3 inhibitor advanced to clinical trials (**Fig.1D-G and Supplementary Fig.3**).

Altogether, these findings established MPiN as a potent and highly selective K_V_1.3 inhibitor with therapeutic potential in autoimmune disease. More importantly, its unprecedented selectivity within the highly conserved K_V_1 subfamily (∼60-213-fold preference for K_V_1.3 over other K_V_1.x channels) provided strong validation of our divergent-pocket targeting strategy and suggests that MPiN engages a previously unrecognized K_V_1.3 binding mechanism distinct from those utilized by conventional, broadly acting K_V_ channel inhibitors.

### MPiN bound a gating-operated pocket in K_V_1.3

The gating of the K_V_1.3 channel is modeled using a simple three-states Markov chain representing sequential transitions between closed (C), open (O), and inactivated (I) states (**Fig.2A,G**). We explored if MPiN state-dependently inhibited the K_V_1.3 channel by measuring channel activity after exposing it to MPiN while stabilized in different gating states. The K_V_1.3 channel was activated using repetitive step depolarizations (20 ms or 2 s) to record stable current, then maintained in the closed state by holding at −90 mV and simultaneously incubated with 10 μM MPiN for 5 min, following which depolarization trains were recovered to analyze channel activity over time (**Supplementary Fig.4A**). It was found that the first current recorded after MPiN incubation showed only minimal inhibition compared to pre-treatment levels; however, currents progressively decreased over subsequent depolarizations, reaching steady-state inhibition faster with 2 s than 20 ms depolarizations (**Fig.2B, Supplementary Fig.4C,E**). Control recordings without compound remained largely stable throughout (**Fig.2B, Supplementary Fig.4B,D**).

**Figure 2.**
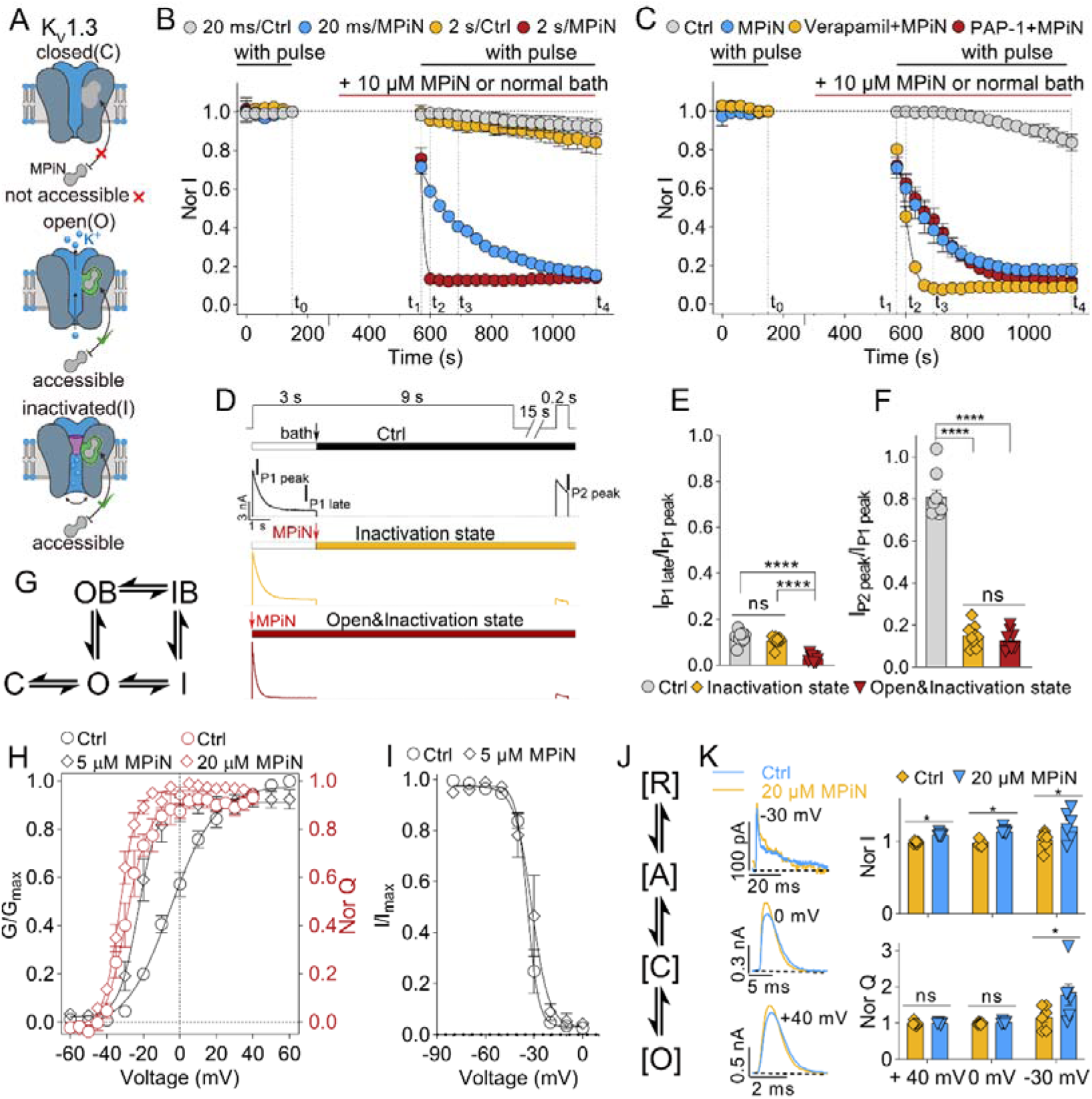
MPiN state-dependently inhibited K_V_1.3 while facilitating its voltage-dependent activation. (A) Schematic depicting MPiN’s accessibility to its binding pocket on K_V_1.3 across the closed, open, and inactivated states. (B) Time course of normalized K_V_1.3 currents (I/I_t0_) during acute perfusion with control bath solution (Ctrl) or 10 μM MPiN (red bar). Channels were activated by +40 mV pulse trains (20 ms or 2 s) during the indicated window (black bar; “with pulse”); otherwise held at −90 mV. n = 8-11. (C) Same protocol as (B), except: (i) pulse duration was 50 ms; (ii) in the “PAP1 + MPiN” and “verapamil + MPiN” groups, channels were preincubated with 200 nM PAP1 or 100 μM verapamil to reach sub-saturated steady-state inhibition before MPiN application (n = 6-13). (D) Representative K_V_1.3 currents evoked by a two-pulse protocol [P1: +40 mV, (3 + 9) s; P2: +40 mV, 0.2 s; 15 s interpulse interval at −90 mV], recorded under control conditions or with MPiN perfusion initiated at the arrowheads (n = 8-9). I_P1 peak_ and I_P2 peak_, peak currents during P1 and P2; I_P1 late_, inactivation-resistant current measured at 3 s of P1. All traces were recorded from the first run of the protocol. (E, F) Summary of I_P1 late_/I_P1 peak_ (E) and I_P2 peak_/I_P1 peak_ (F) ratios from (D), representing the fractions of inactivation-resistant and recovered channels, respectively (n = 8-9). (G) Kinetic scheme of K_V_1.3 gating transitions among closed (C), open (O), and inactivated (I) states. MPiN binds the open or inactivated channel, trapping it in the open-blocked (OB) or inactivated-blocked (IB) state. (H, I) Normalized conductance-voltage (G-V; black) and gating charge-voltage (Q-V; red) relationships (H), and steady-state inactivation curves (I) of K_V_1.3 in the presence or absence of MPiN (n = 6-12). (J) Allosteric model of K_V_1.3 activation. R↔A, voltage sensor transition from resting to activated; C↔O, pore transition from closed to open; J, allosteric coupling factor. (K) Left: representative outward gating currents of K_V_1.3 evoked at various depolarizing voltages with or without MPiN. Right: normalized gating current (upper) and gating charge (lower), each scaled to the mean control value at the corresponding voltage (n = 6).

Verapamil is an antiarrhythmic drug that inhibits K_V_1.3 by targeting the channels’ open and inactivated states. It traps channels in an inactivated blocked state, from which they recovered much more slowly than from the drug-free inactivated state^39^. We reasoned that this property would provide extended opportunity for MPiN binding and accelerate its inhibition kinetics. This hypothesis was tested using a slightly modified protocol with 50-ms depolarizations (**Supplementary Fig.4F**). Indeed, verapamil-treated K_V_1.3 channels were not further inhibited significantly over time, but were inhibited much faster by MPiN than untreated channels (**Fig.2C, Supplementary Fig.4G-I**). In contrast, another K_V_1.3 inhibitor, PAP-1, did not alter MPiN inhibition kinetics (**Fig.2C, Supplementary Fig.4J**). These findings demonstrated that MPiN did not target closed-state K_V_1.3 channels; rather, inhibition occurred after channel opening, potentially involving open-state binding, inactivated-state binding, or both.

The fact that MPiN effectively inhibited K_V_1.3 currents during 20 ms depolarizations, which induced minimal C-type inactivation, suggesting that it could inhibit K_V_1.3 channels through open-state interaction alone (**Fig.2B, Supplementary Fig.4C**). We subsequently examined MPiN’s ability to interact and block the inactivated channels. The voltage-clamp and compound perfusion protocols are as shown in Figure 2D. Channels were activated and driven into a stable inactivated state using the first +40 mV/12 s depolarization pulse (P1), and those channels recovered during a 15 s holding at −90 mV were measured using the second +40 mV/0.2 s test pulse (P2). MPiN was applied at 3 s of the test pulse P1 to specifically target channels predominantly in their inactivated state, or in a preincubation manner (before pulse) to broadly target channels in their open and inactivated states. The peak currents at test pulses P1 and P2, and the inactivation-resistant residual current at 3 s of P1, were defined as I_P1 peak_, I_P2 peak_, and I_P1 late_, respectively (**Fig.2D)**. In the absence of MPiN, the majority of channels transitioned into the inactivated state after a 3 s depolarization, with a I_P1 late_/I_P1 peak_ ratio of 12.1 ± 0.8% (**Fig.2D, E**). These inactivated channels were largely recovered after 15 s repriming at −90 mV, as demonstrated by a I_P2 peak_/I_P1 peak_ ratio of 80.8 ± 3.1% (**Fig.2D, F**). MPiN applied at 3 s of the test pulse P1 significantly reduced the I_P2 peak_/I_P1 peak_ ratio to 14.9 ± 1.6%, without markedly affecting the I_P1 late_/I_P1 peak_ ratio (10.7 ± 0.7%); moreover, preincubated MPiN significantly reduced both the I_P2 peak_/I_P1 peak_ and I_P1 late_/I_P1 peak_ ratios to 12.8 ± 1.6% and 2.9 ± 0.5%, respectively (**Fig.2D-F**). These data provided strong evidence that MPiN effectively targeted channels in the inactivated state.

Collectively, our findings demonstrated that MPiN acted on post-opening K_V_1.3 channels, trapping them in the open blocked (OB) and inactivated blocked (IB) states (**Fig.2G**), by associating with a binding pocket operated by the channel’s gating process, in which a closed conformation transition buries the binding pocket, whereas an open and inactivation conformation transition exposes it (**Fig.2A**).

### MPiN uniquely facilitated K_V_1.3 activation gating

Mechanistically, MPiN inhibited K_V_1.3 either by modulating channel gating or physically blocking channel pore, two mechanisms that are expected to produce distinct effects on gating kinetics. MPiN voltage-independently inhibited K_V_1.3 currents across a wide range of depolarizing voltages (**Supplementary Fig.5A-C**). Unexpectedly, however, it simultaneously facilitated the channel’s voltage-dependent activation, producing a significant hyperpolarizing shift of the steady-state activation curve (**Fig.2H, black curves; Supplementary Fig.5D**), while leaving steady-state inactivation unaffected (**Fig.2I, Supplementary Fig.5E**). This unusual behavior resembles the gating-modifying action of β-scorpion toxins on Na_V_ channels ^40^.

Characteristic of voltage-gated ion channels, K_V_1.3 activation involves three coordinated processes: voltage sensor activation (R↔A), pore opening (C↔O), and their electromechanical coupling (**Fig.2J**). To identify MPiN’s action within this cascade, we directly measured its effect on voltage sensor activation by measuring gating currents. MPiN significantly increased gating currents amplitude at weak (−30 mV), moderate (0 mV), and strong (+40 mV) depolarizations, indicating enhanced synchronization of voltage sensor activation by the compound (**Fig.2K, left and right upper panels**). However, the total gating charges transferred, quantified by integrating gating current over time, was increased only during weak depolarization (−30 mV) (**Fig.2K, right lower panel**). Consistent with these observations, MPiN induced a hyperpolarizing shift in the voltage sensor activation curve, promoting voltage sensor activation at voltages below −15 mV (**Fig.2H, red curves; Supplementary Fig.5F-G**). These data confirmed that MPiN facilitated voltage sensor activation at weak depolarizations, with this effect being progressively overcome at stronger depolarizations. Moreover, it was noteworthy that MPiN facilitated K_V_1.3 channel activation without affecting its gating charge activation at 0 mV, suggesting that the compound also acted on the downstream pathway of voltage sensor activation (**Fig.2H**). On the other hand, MPiN did not alter the time-dependent activation and deactivation of K_V_1.3, implying it modified the energetic landscape of voltage sensor activation without affecting the intrinsic transition rates between distinct conformational states (**Supplementary Fig.5H**). To our knowledge, MPiN was the first K_V_1.3 inhibitor to both inhibit channel activity while facilitating channel activation, suggesting a distinct action mechanism from documented K_V_1.3 inhibitors.

### Two K_V_1.3-specific residues jointly determined selective MPiN recognition

To map the molecular determinants on K_V_1.3 determining selective MPiN interaction, we first examined whether MPiN competed with reported K_V_1.3 inhibitors of distinct action mechanisms, including the peptidic outer pore blockers ShK and ADWX-1, the small molecule inner pore blockers PAP-1 and verapamil, and TEA-Cl which acts on both sides of the pore (**Fig.3A**). To maximize binding-site occupancy and enable robust competition with MPiN, these competing inhibitors were applied by preincubating with K_V_1.3 channels at high concentrations, producing 68.0-91.1% currents inhibition (**Fig.3B, Supplementary Fig.6A-F**). Results showed that ADWX-1, but not the others, significantly impaired MPiN’s inhibitory potency on K_V_1.3, by right forward shifting the dose-response relationship, raising the apparent IC_50_ value by approximately 10-fold (**Fig.3C, Supplementary Fig.6G-H**). Preincubation with 10 μM MPiN drastically inhibited K_V_1.3 currents and reversibly reduced ADWX-1’s inhibitory potency by approximately 3-fold (**Fig.3D, Supplementary Fig.6I-K**). Despite the observed asymmetric competition effect, these data provided strong evidence supporting an interaction between ADWX-1 and MPiN in their action with K_V_1.3. It was likely that MPiN targeted the S5-S6 loop in K_V_1.3 as ADWX-1. Sequence alignment of MPiN-sensitive K_V_1.3 with other MPiN-resistant K_V_1.x channels revealed two short K_V_1.3-specific sequences- “_423_DPTSG_427_” located within the PP1 linker connecting S5 to the pore helix, and “_451_HPVTI_455_” within the PP2 linker connecting the selectivity filter to S6, in the S5-S6 region (**Supplementary Fig.7**). Indeed, the chimeric channel constructed by replacing the K_V_1.5 PP1 and PP2 linkers with their K_V_1.3 counterparts (K_V_1.5/1.3-P1P2) gained robust MPiN sensitivity, comparable to K_V_1.3, highlighting critical roles of these two variable sequences in selective MPiN recognition (**Fig.3E,F, Supplementary Fig.8**). Based on these findings, scanning mutagenesis of K_V_1.3 residues to its counterpart in K_V_1.5 further identified two key residues: the 427^th^ glycine (G427) in the PP1 linker and the 451^th^ histidine (H451) in the PP2 linker, with G427H and H451R mutations reducing MPiN potency by 158.4-fold and 94.2-fold, respectively (**Supplementary Fig.9A, B, E**). Regarding the K_V_1.5 reverse mutants, while single-site reverse mutation of the 463^th^ histidine or the 487^th^ arginine in K_V_1.5 to their counterparts in K_V_1.3 respectively (H463G or R487H) failed to confer MPiN sensitivity, the double mutation H463G/R487H fully restored inhibition to levels comparable with K_V_1.3 (**Fig.3E-F, Supplementary Fig.8, Supplementary Fig.9C-E**). Mutating other variable residues in K_V_1.5 S5-6 loop to their corresponding residues in K_V_1.3 did not enhance MPiN’s inhibitory potency (**Supplementary Fig.9C-E**). Accordingly, our data demonstrated that the G427 and H451 residues in K_V_1.3 exclusively determined selective MPiN recognition. Notably, these two residues also jointly underlay ADWX-1’s high selectivity for K_V_1.3 over K_V_1.5 (**Fig.3F, Supplementary Fig.10A-C**). Compared with MPiN, ADWX-1 had distinct effects on the gating of K_V_1.3 by left forwardly shifting the channel’s steady-state inactivation relationship without affecting the channel’s voltage-dependent activation (**Supplementary Fig.10E-I**). These data, together with our in-depth mutation analysis addressed below, demonstrated that MPiN and ADWX-1 utilized the same molecular machinery, comprised of the K_V_1.3-specific G427 and H451 residues, to selectively inhibit the K_V_1.3 channel, but by targeting different binding pockets on the channel.

**Figure 3.**
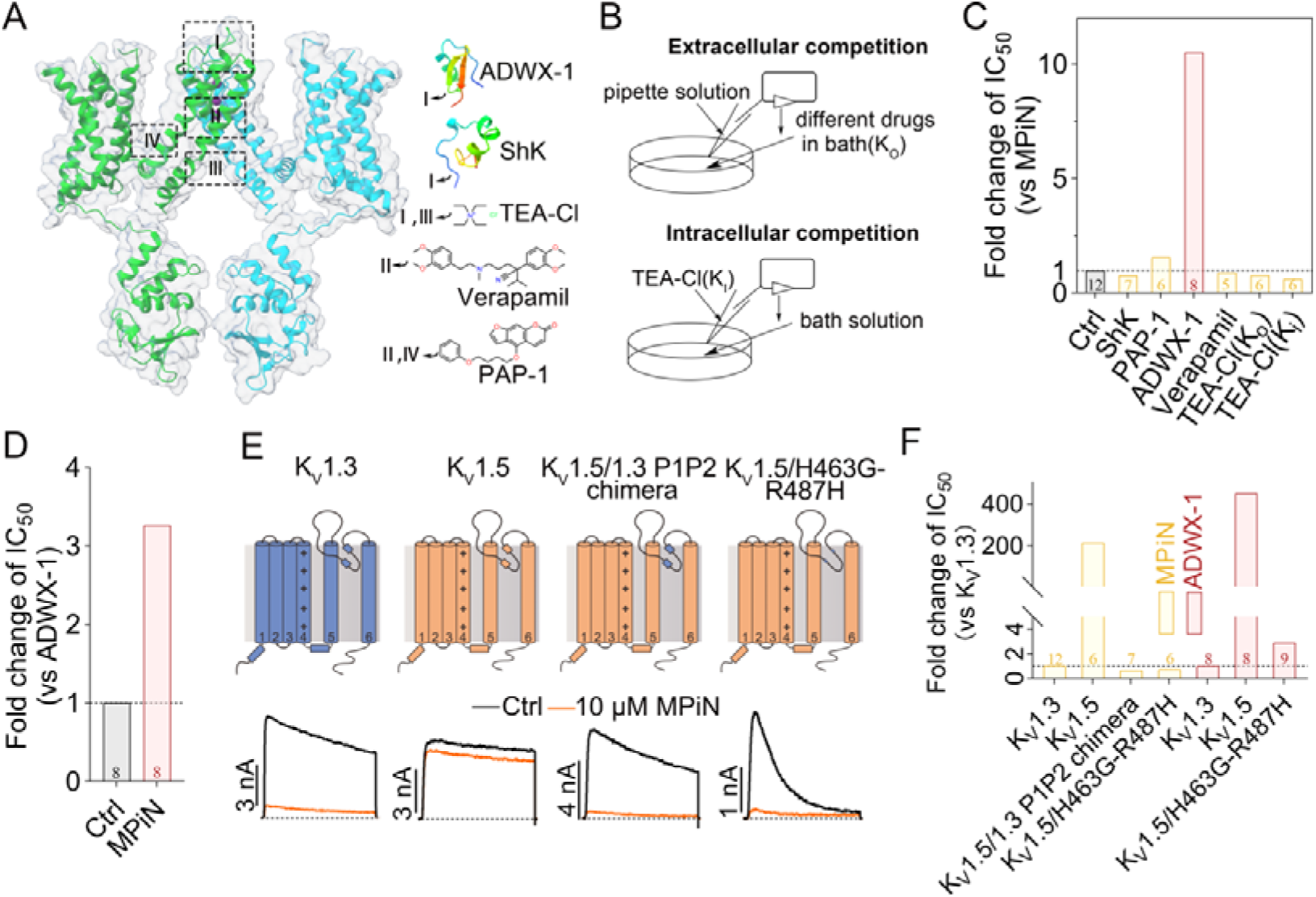
Molecular determinants of selective MPiN recognition by K_V_1.3. (A) Binding sites of previously reported K_V_1.3 inhibitors mapped onto the channel structure. (B) Schematic of the drug competition assay between known K_V_1.3 inhibitors and MPiN. Cells were preincubated with ShK, PAP-1, ADWX-1, or verapamil (in bath), or with TEA-Cl (in bath or pipette), before MPiN was acutely perfused. (C)-(D) Fold change in IC_50_ of MPiN (C) or ADWX-1 (D) inhibiting K_V_1.3 in the presence of each preincubated competing drug as indicated, relative to the IC_50_ measured without preincubation (control) (n = 5-8).(E) Topological diagrams of the K_V_1.3, K_V_1.5, the K_V_1.5/1.3 P1P1 chimera, and K_V_1.5-H463G/R487H mutant channels, with their representative currents before and after application of 10 μM MPiN (n = 6-9). (F) Fold change in IC_50_s of MPiN (yellow) and ADWX-1(pink) inhibiting K_V_1.5, K_V_1.5/1.3 P1P1 chimera, and K_V_1.5-H463G/R487H mutant channels, relative to their respective IC_50_s of inhibiting K_V_1.3 (n = 6 - 9).

### A novel inhibitor binding pocket in K_V_1.3 probed by MPiN

To define the MPiN binding pocket in K_V_1.3, we performed alanine/cysteine-scanning mutagenesis across the transmembrane S1-6 helices and the S4-5 and S5-6 loops, encompassing all regions potentially accessible to small molecule inhibitors. Multiple mutations in the S2, S3, S5, and S6 helices and the extracellular S5-6 loop (P-loop) markedly reduced MPiN potency (**Supplementary Fig.11**). The apparent IC_50_ values as determined by concentration-response relationships were increased by 5 to 10-fold for the F306A, I402A,V409A, G456A, L474A, 10-20 fold for the F403A, A419C, A434C, S462A, and > 25 fold for the V416A, D433A, F435A, A465C mutations (**Fig.4A, upper panel; Supplementary Fig.12A-C**). Mapping these key residues together with G427 and H451 onto the K_V_1.3 structure revealed a contiguous cluster surrounding a pocket formed by the extracellular ends of the S5 and S6 helices, the PP1 and PP2 linkers, and the pore helix (**Fig.4B**). Notably, the most influential residues, including G427, H451, D433, and F435, were located near the pocket center (**Fig.4B**). In contrast to D433, other MPiN-sensitive residues were dispensiable for ADWX-1 activity (**Fig.4A, lower panel; Supplementary Fig. 12D,E**). Consistent with D433 serving as the sole shared interaction site, the D443A mutation completely abolished the competition between MPiN and ADWX-1 (**Fig.4C, Supplementary Fig.13A-C**). These findings indicated that MPiN engaged a binding pocket distinct from that of ADWX-1, while both ligands exploited the same K_V_1.3 selectivity determinants, G427 and H451.

**Figure 4.**
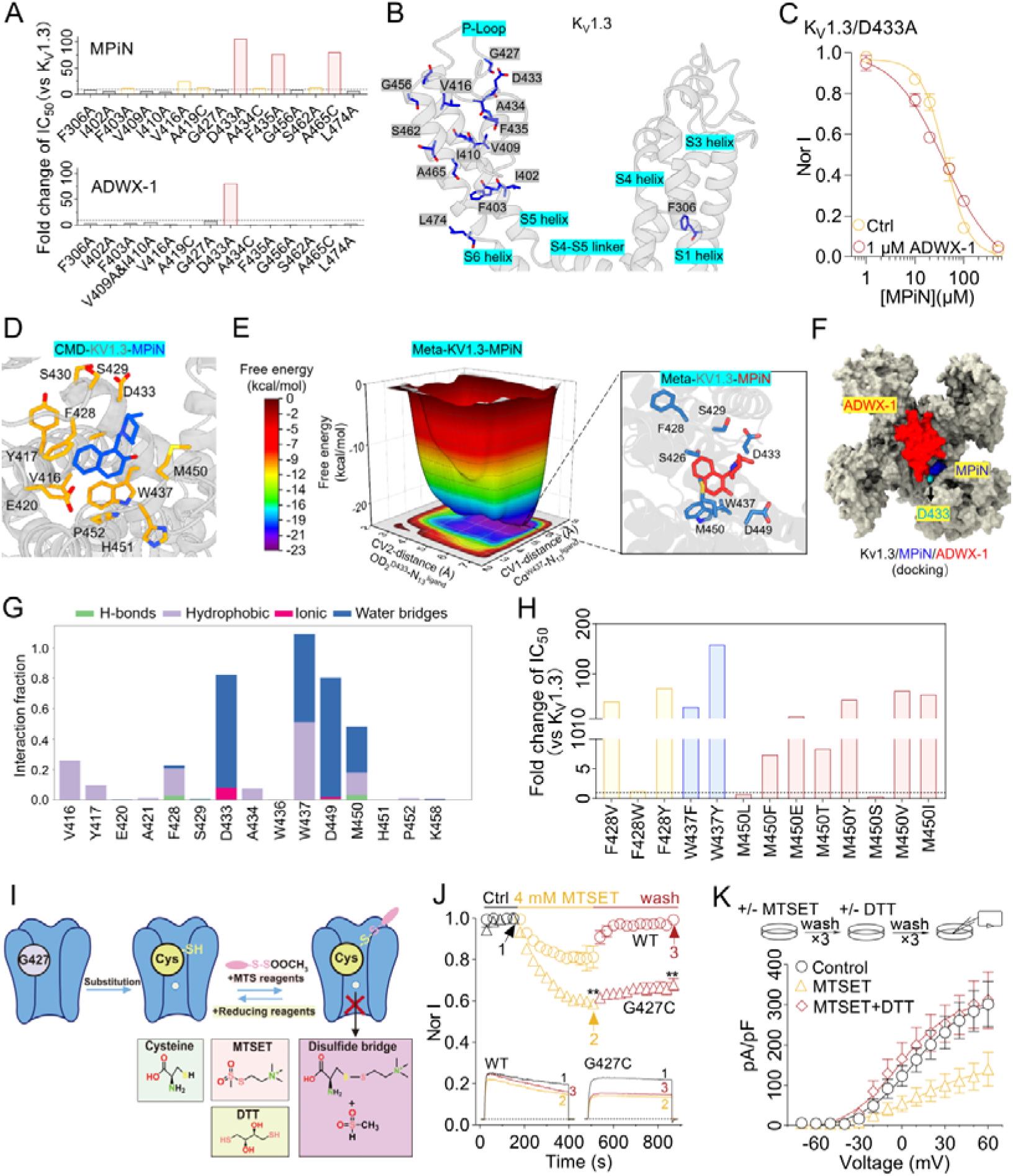
Identification of a novel MPiN-K_V_1.3 binding pocket by molecular dynamics and large-scale mutagenesis. (A) Comparative alanine-scanning mutagenesis reveals overlapping yet distinct interaction patterns of MPiN and ADWX-1 with K_V_1.3. The D433A mutation significantly reduces the IC_50_ values of both MPiN (top) and ADWX-1 (bottom) (n = 5-8). (B) K_V_1.3 structure (gray) with alanine-scanning mutation sites depicted as blue sticks. (C) Concentration-response curves for competitive inhibition of K_V_1.3-D433A by ADWX-1 and MPiN (n = 5-6). (D) Ligand-protein interaction interface between MPiN and K_V_1.3. Ligands are shown as blue sticks; key residues as yellow sticks; the receptor is rendered as a gray cartoon. (E) Lowest-energy conformations of MPiN and K_V_1.3 identified by MetaD simulations.(F) Structural superposition showing ADWX-1 (red surface) and MPiN (blue surface) bound to K_V_1.3 (gray surface). The overlapping residue D433 is highlighted as a cyan solid surface. (G) Bar chart of binding interaction forces between MPiN and K_V_1.3 during conventional molecular dynamics (CMD) simulations. (H) Fold change in IC_50_ for MPiN inhibition of K_V_1.3 mutants relative to wild-type (WT), as determined by electrophysiological recordings (n = 5-7). All data are presented as mean± SEM (n = 5-7). (I) Schematic diagram of the MTSET covalent modification strategy. (J) Time-course of K_V_1.3 and K_V_1.3/G427C currents inhibition by acute MTSET (4 mM) perfusion and subsequent recovery upon washout, inset shows representative traces at indicated time points (n = 4-6). (K) Upper, treatment protocol: channels were modified with MTSET (4 mM, 10 min), washed three times with bath solution, then treated with DTT (4 mM, 10 min) or left untreated (± DTT); controls received neither reagent (−MTSET/-DTT). Each reagent step was followed by three bath-solution washes prior to patch-clamp recording. Lower, current-voltage (I-V) relationships of K_V_1.3-G427C under the indicated conditions (n = 10-13).

To resolve the binding modes at atomic resolution, MPiN and ADWX-1 were subjected to induced-fit docking into the C-type conformational model of K_V_1.3, followed by all-atom conventional molecular dynamics (CMD), metadynamics (MetaD) simulations, and mutagenesis-based functional validation. The binding poses obtained from CMD and MetaD simulations were highly consistent for both ligands (**Fig.4D,E; Supplementary Fig.14A-C**). ADWX1 adopted the characteristic pore-blocking binding mode, occupying the extracellular entrance of the pore and forming extensive polar interactions with surrounding residues (**Supplementary Fig.14B-D**). K26 inserted into the pore center and formed a stable hydrogen bond with T446 (**Supplementary Fig.14D**). In contrast, MPiN bound within the central region of the S5-S6 loop, with its hydrophobic heterocyclic moiety deeply embedded between the S5 and S6 helices (**Fig.4D,E**). This binding mode was stabilized by hydrophobic contacts with V416, Y417, F428, W437, and M450, whereas its hydroxyl and imine groups formed persistent salt-bridge and water-mediated interactions with D433 and D449 (**Fig. 4G**).

Notably, K8 and R23 of ADWX1 formed salt bridges with D433 on chains A and C of K_V_1.3, respectively. This interaction site coincided with the D433 residue involved in MPiN binding (**Fig. 4F, Supplementary Fig.14D**). These computational findings are consistent with the competitive binding and mutagenesis data (**Fig.3C,D and Fig.4C**), supporting the proposed binding modes of MPiN and ADWX1 on K_V_1.3.

To further validate the proposed MPiN-binding pocket, mutations were introduced at residues F428, W437, and M450 identified by CMD and MetaD simulations (**Fig.4G**), and the effects on the IC_50_ for MPiN-mediated inhibition of K_V_1.3 activation were determined (**Supplementary Fig.15**). Mutations F428V, F428Y, W437F, W437Y, M450E, M450Y, M450V, and M450I reduced the apparent affinity of MPiN by approximately 10-200-fold (**Fig.4H**), whereas M450T, M450F, M450L, and M450S caused only modest changes in apparent affinity (**Fig.4H**).

In addition, a covalent occupancy strategy^41,42^ was employed to independently verify MPiN binding (**Fig.4I**). In the Kv1.3-G427C mutant, treatment with 4 mM MTSET significantly reduced the inhibitory effect of MPiN on Kv1.3 currents, whereas subsequent treatment with DTT restored MPiN-mediated inhibition (**Fig.4J,K**). Despite the absence of a direct interaction between G427 and MPiN, the position of G427 within the binding pocket and the structural effects of covalent occupancy at C427 support the existence of an MPiN-binding site. These findings further support the conclusion that MPiN binds within the cleft formed by the S5-S6 loop and the S5 and S6 helices of K_V_1.3.

### The selectivity-determining residues G427 and H451 maintained the structural integrity of MPiN binding pocket in K_V_1.3

We next determined the structural basis by which G427 and H451 confer selective recognition of MPiN by K_V_1.3. MetaD simulations of the K_V_1.3/G427H mutant showed that the introduced H427 residue on the P1 loop approached H451 on the P2 loop, resulting in a marked reduction in the distance between the two loops (**Fig.5A-D**). This conformational change narrowed the inter-loop cleft and hindered insertion of MPiN into the binding pocket. In addition, the larger histidine side chain likely introduced steric hindrance that further reduced ligand binding (**Fig.5C**).

**Figure 5.**
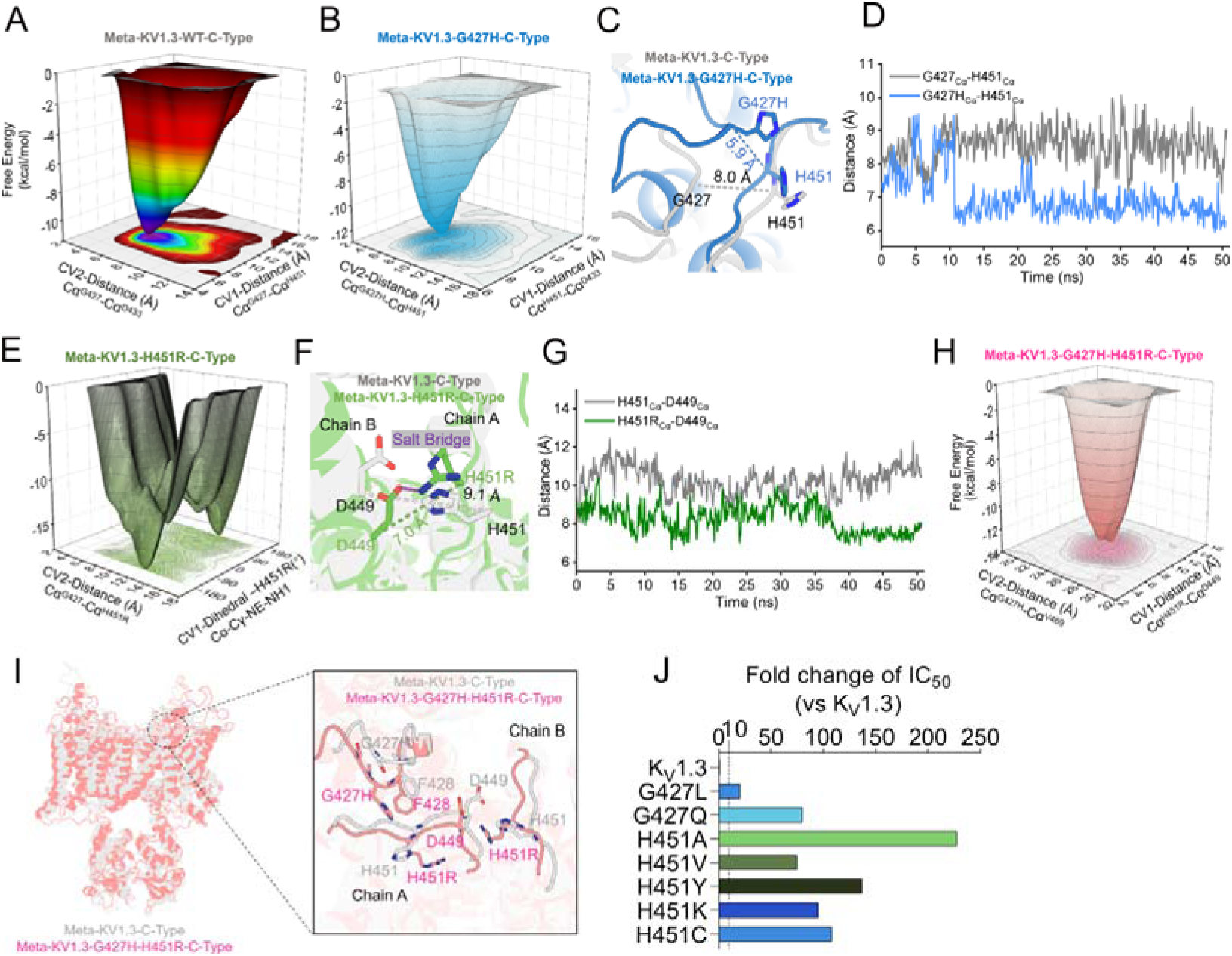
Structural basis of G427 and H451 in determining selective MPiN recognition in K_V_1.3. (A-I) Conformational changes in the P1 and P2 loops of Kv1.3-C-type (A) and the G427H (B-D), H451R (E-G), and G427H-H451R (H,I) mutants revealed by metadynamics (MetaD) simulations. (J) Fold change in MPiN IC_50_ for indicated mutant channels relative to wild-type K_V_1.3 (n = 6-7).

MetaD simulations of the Kv1.3/H451R mutant showed that the H451R substitution forms a strong electrostatic interaction with D449, reducing the distance between the P2 loop of chain A and chain B and inducing a conformational change in the P2 loop (**Fig.5A,E-G**). This change is likely to reduce the stability of MPiN binding. In addition, MetaD simulations of the Kv1.3G427H/H451R double mutant revealed conformational changes in both the P1 and P2 loops (**Fig.5H,I**).

To validate the MetaD models and clarify the mechanisms of MPiN resistance observed in other K_V_1.x channels besides K_V_1.5, we conducted functional mutagenesis analysis through mutating G427 and H451 to their corresponding residues in K_V_1.1-1.2, K_V_1.4, and K_V_1.6-1.8. It was found that mutations G427A, G427L, and G427Q significantly increased the IC_50_ of MPiN (**Fig.5J, Supplementary Figure 16**), and the magnitude of the increase correlated with the size of the substituted side chain (**Fig.5E**), supporting the computational model. Mutations at H451 (H451A, H451V, H451Y, H451K, and H451C) similarly increased the IC_50_ of MPiN by approximately 50–250 - fold (**Fig. 5E, Supplementary Figure 16**).

Together, these results argued that G427 in the PP1 loop and H451 in the PP2 loop contribute to maintaining the structural integrity of the MPiN binding pocket in K_V_1.3. MPiN achieves subtype selectivity through recognition of these two non-conserved residues.

### MPiN allosterically inhibited the K_V_1.3 channel

We investigated the mechanism by which MPiN allosterically inhibits K_V_1.3 activation, which involves coordinated conformational coupling among the PP1 and PP2 loops and the S5/S6 helices, thereby restricting K^+^ permeation across the membrane (**Fig.6A**). Structural alignment of the CMD-derived K_V_1.3 C-type state with the MPiN-bound, CMD-refined complex shows that MPiN disrupts the π–π stacking interaction between F428 and W437, increases the distance between the PP1 and PP2 loops, and shifts the PP2 loop toward the voltage-sensing domain (VSD) (**Fig.6A, left**). These changes propagate to the S5 helix, S6 helix, and pore helix, leading to rearrangement of residues V416, F435, W436, V410, V409, and V439 (**Fig.6A, middle**).

**Figure 6.**
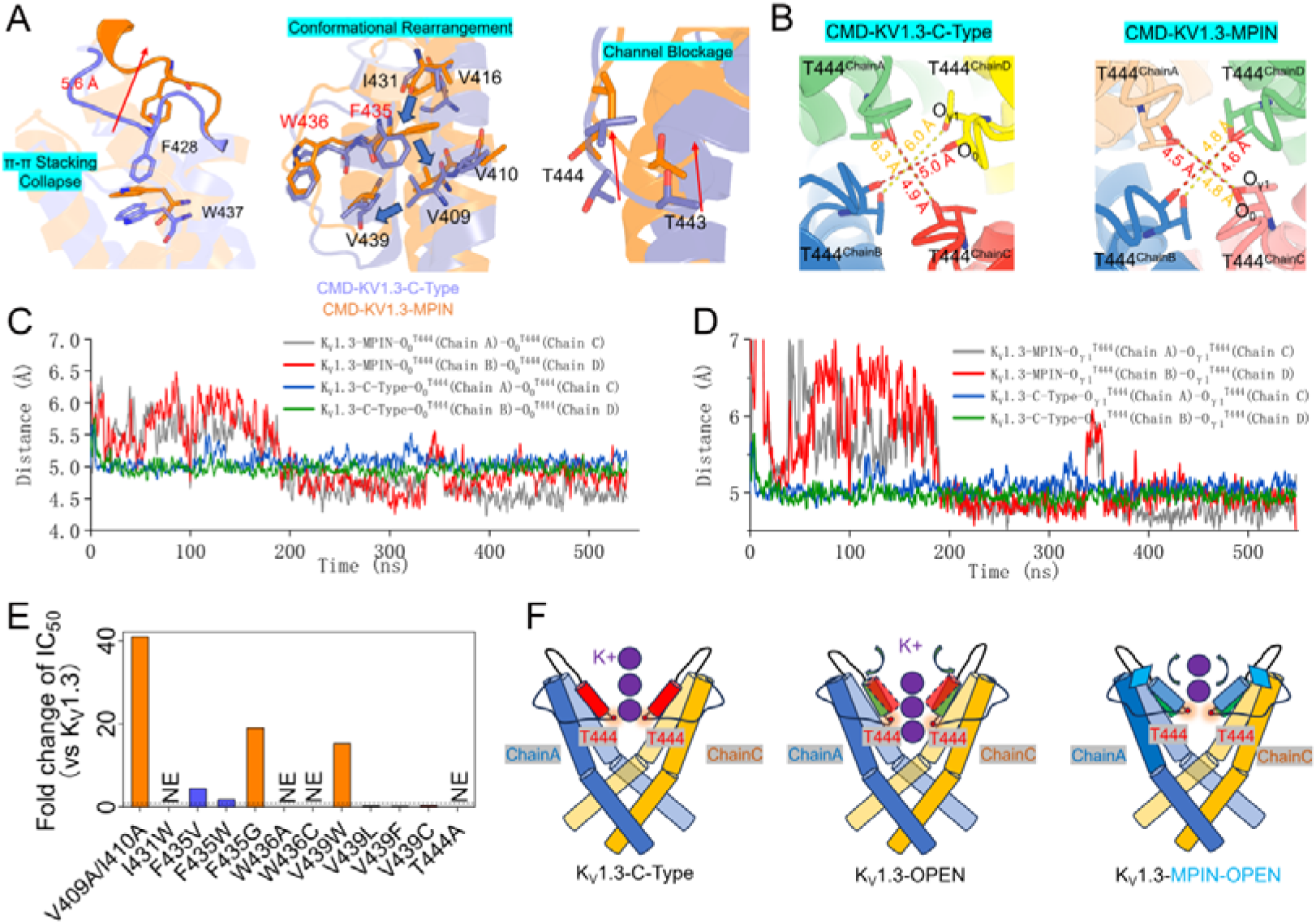
Molecular mechanism of MPiN allosteric inhibition of K_V_1.3 activation. (A) CMD simulations reveal the conformational propagation pathway induced by MPiN binding. (B-D) Inter-chain distances of the equivalent T444 atoms (Oγ1–Oγ1 and O0–O0) in Kv1.3-C-type and Kv1.3-MPiN, shown as structural views (B) and time-dependent distance plots (C, D) derived from CMD simulations. (E) Bar plot of the fold change in apparent ICLL of MPiN for mutants of key residues along the conformational propagation pathway (n = 6-10). (F) Schematic model of the molecular mechanism underlying MPiN-mediated allosteric inhibition of Kv1.3. All data are presented as mean ± SEM (n= 6-10).

V416, V409, and V410 move toward the VSD, while F435 rotates toward S6 and the pore helix shifts upward (**Fig.6A, middle**). These changes bring the P2 loops closer together, resulting in a reduction in pore diameter (**Fig.6A,B**). Distance analysis of the O_γ1_ and O_0_ atoms of T444 indicates that MPiN reduces the K_V_1.3 pore diameter by approximately 0.5 Å (**Fig.6B-D**), stabilizing a more closed state and thereby impeding K^+^ permeation.

Site-directed mutagenesis of residues along the proposed MPiN-induced conformational pathway further supports the CMD observations. The V416A mutation significantly decreased the IC_50_ of MPiN (**Fig. 4A**). Mutations at F435 (F435W and F435V) had little effect on MPiN-induced conformational propagation, whereas F435A and F435G reduced the IC_50_ by more than 50- and 19-fold (**Fig.4A**, **Fig.6E, Supplementary Figure 17**). The V409A/I410A double mutant also reduced IC_50_ by more than 40-fold, and V439W similarly decreased IC_50_ (**Fig.6E, Supplementary Figure 17**). Interestingly, the V439L, V439F, and V439C mutations slightly increased channel sensitivity to MPiN, likely by facilitating the conformation propagation of channel inhibition by the compound (**Fig.6E, Supplementary Figure 17**).

These results support an allosteric pathway from the PP1-PP2 loop through the S6 helix to the pore helix. During voltage activation of K_V_1.3, outward movement of the pore helix expands the pore and enables K^+^ permeation (**Fig.6F**). In contrast, MPiN binds above the pore helix and restricts coupling between the selectivity filter and S6 helix, preventing outward motion of the pore helix, reducing pore radius, and inhibiting K^+^ flux (**Fig.6F**).

## Discussion

Through a rational subtractive screening strategy combined with electrophysiological and computational analyses, we identified MPiN, a novel small-molecule inhibitor with high selectivity for the K_V_1.3 channel. MPiN engages a previously uncharacterized druggable pocket formed by the turret loops, pore helix, selectivity filter, and the extracellular ends of the S5 and S6 helices - a region that is highly conserved across the K_V_1.x subfamily. Binding at this site allosterically couples the pore domain to the voltage-sensing apparatus, trapping the channel in its inactivated conformation and thereby suppressing activation. Remarkably, despite the conservation of the binding pocket itself, MPiN achieves stringent selectivity for K_V_1.3 over other K_V_1.x isoforms through two peripheral residues, G427 and H451, that shape the pocket without directly contacting the ligand. In a murine psoriasis model, MPiN exhibited therapeutic efficacy comparable to the clinical candidate PAP-1, establishing it as a viable template for next-generation K_V_1.3 drug design. Collectively, our findings define a new paradigm for the selective pharmacological control of K_V_1.3 and uncover a previously unrecognized druggable pocket, providing a structural foundation for the rational design of a new generation of K_V_1.3-targeting therapeutics.

Among K_V_1.3-selective inhibitors in clinical development, DES7114 lacks publicly disclosed mechanistic data^38^, leaving PAP-1 - derived from the psoralen scaffold Psora-4 - as the only high-selectivity K_V_1.3 blocker with a well-defined mode of action^43^. Psora-4 inhibits K_V_1.5 via a dual-site mechanism, co-occupying the conserved central pore cavity and an allosteric “side pocket” formed at the inter-subunit interface by S4, the S4-S5 linker, S5, S6, and the pore helix; cooperative binding (Hill≈2.8) drives an inactivation-like filter collapse, producing nanomolar potency (IC_50_≈3 - 7 nM), strong use-dependence, and very slow recovery, with subtype selectivity dictated almost entirely by the non-conserved side pocket^33^. Consistently, PAP-1 inhibits K_V_1.3 at IC_50_≈2 nM but shows only ∼23-fold selectivity over K_V_1.5. MPiN operates through a fundamentally different paradigm. Despite four equivalent binding sites on the tetramer, its Hill coefficient of ∼1.26 reflects a 1:1, non-cooperative interaction; more remarkably, its K_V_1.3 selectivity is conferred entirely by two peripheral, non-contact residues - G427 and H451 - that act through long-range allostery. This mirrors the Na_V_1.8-selective inhibitor A - 803467, whose selectivity is likewise dictated by two non-contact residues (Thr397, Gly1406) in an otherwise conserved vestibule^44^, hinting at a generalizable “non-contact selectivity” paradigm. Although less potent than PAP-1, MPiN exhibits superior intra-K_V_1.x selectivity and matches PAP-1’s efficacy in a mouse psoriasis model. As the first entirely new pharmacological site identified on K_V_1.3 in nearly three decades, the MPiN pocket provides a prototype scaffold for next-generation K_V_1.3 drug design. This finding parallels recent discoveries of unconventional allosteric sites across the K_V_ family - the AUT5/AUT1-induced extracellular “turret cap” at the K_V_3 VSD-PD interface ^45^, the spatially distinct transmembrane VSD-PD pocket of Lu AG00563^46^, and the pore-domain lateral fenestration engaged by retigabine in K_V_7^47^ - collectively revealing an expansive, largely untapped landscape of druggable K_V_ allosteric sites that, in the era of AIDD and CADD, offer a timely foundation for structure-guided drug design.

Certain voltage-gated K^+^ channel blockers exhibit a paradoxical dual action: while inhibiting the macroscopic current, they simultaneously shift the activation curve toward more hyperpolarized potentials through allosteric coupling to the voltage-sensing domain or its interface with the pore^48–50^. Here we show that MPiN, at sub-blocking concentrations, markedly enhances the voltage sensitivity of K_V_1.3, left - shifting V_a_ and reducing the slope factor, whereas the pore-blocker ADWX1 is devoid of this effect. This divergence reflects distinct binding modes. ADWX1 docks directly above the selectivity filter, physically obstructing ion flux without engaging the VSD, and thus leaves gating kinetics unperturbed. MPiN, in contrast, inserts into a recessed pocket between the PP1 and PP2 loops, adjacent to S6 and the apex of the PP2 helix. Steric engagement at this site drives an outward displacement of S6 and the PP2-helix apex, shortening the distance between S6 and core VSD residues. We propose that this rearrangement reinforces the R364(S4)-D422(S6) salt bridge, thereby accelerating gating - charge translocation and stabilizing the activated VSD, which lowers the energetic barrier for pore opening (**Supplementary Figure 18**). MPiN therefore represents a bifunctional modulator that simultaneously blocks ionic conductance and allosterically potentiates VSD activation, while ADWX1 behaves as a pure pore blocker. Beyond mechanistic interest, this contrast offers a chemical tool to dissect the allosteric coupling network linking the pore domain to the voltage sensor in K_V_1.3.

MPiN inhibits K_V_1.3 in a strictly state-dependent manner, engaging only channels in the open or inactivated conformation - a consequence of the binding pocket being exposed exclusively in these two states^1,6,32^. Such strict state-dependence is a shared feature among a panel of K_V_-channel modulators, though the underlying mechanisms are diverse. The classical blocker TEA enters the inner pore cavity from the cytoplasmic side and both occludes K^+^ permeation and impedes activation-gate closure via the canonical “foot-in-the-door” mechanism, slowing channel deactivation^51,52^ Verapamil binds the central cavity at A413/I420, preferentially targets the open and inactivated states, and drives the drug-bound channel into a slowly recovering “inactivated-blocked” state^28,53^. Clofazimine exhibits a dual-mode block - open-channel block combined with lock-in of the channel in a “closed-deactivated” state - with both modes requiring prior channel opening and sparing resting closed and C-type inactivated channels^27^. Despite these mechanistic differences, state-dependent inhibitors converge on a shared advantage analogous to a “servo” mechanism: they act only when pathologically relevant channels are engaged, potentially mitigating off-target effects from over-inhibition - a paradigm to which MPiN’s stringent state-dependence firmly conforms, adding a meaningful developability advantage to its next-generation K_V_1.3-targeted profile.

In summary, MPiN stands as a first-in-class K_V_1.3 chemotype of exceptional therapeutic potential, and the previously hidden, selectivity-encoding druggable pocket it unveils opens a new chapter in structure-guided drug design against the voltage-gated ion-channel superfamily.

## Materials and methods

### Compounds screening

A structurally diverse compound library comprising 4,208 compounds was purchased from Selleck Chemicals (Catalog No. L3600; Selleckchem, Houston, TX). The K_V_1.3 channel was heterologously expressed in HEK293T cells, and whole-cell currents were recorded using patch-clamp electrophysiology with +20 mV/300 ms test pulses from a holding potential of −80 mV at 5-s sweep intervals. After stable K_V_1.3 currents were established, normal bath solution was applied as an internal control, followed by administration of test compounds (10 μM), both via acute perfusion, and the extent of current inhibition was recorded. In most cases, the compounds had little or no effect on K_V_1.3 currents; therefore, up to five compounds were tested sequentially in a single recording cell, after which TEA-Cl was administered to fully inhibit the currents and the cell was discarded. Each compound was tested in three independent recordings.

### Drugs

MPiN, the hit compound in the screening analysis, was synthesized by SUSIMM (Suzhou Institute of Materia Medica, Suzhou, China), and its structure and purity were validated by quality-control analyses using nuclear magnetic resonance (NMR), high-performance liquid chromatography (HPLC), and electrospray ionization mass spectrometry (ESI-MS). PAP-1 and verapamil were purchased from Aladdin Biochemical Technology, and TEA-Cl was obtained from Sigma-Aldrich. Stock solutions (100 mM) were prepared in DMSO and stored at −80 °C. Working solutions were freshly prepared by diluting stock solutions at least 200-fold into the appropriate extracellular solution, resulting in a final DMSO concentration of <0.5%. Linear ADWX-1 was synthesized by Fmoc-based solid-phase peptide synthesis. The purified reduced peptide was refolded at 4L for 16 h in a buffer containing 0.1 M Tris, 0.1 M NaCl, 5 mM reduced glutathione (GSH), 0.5 mM oxidized glutathione (GSSG), and 2 mM EDTA (pH 9.3, adjusted with NaOH). The refolded peptide was purified by reverse-phase high-performance liquid chromatography (RP-HPLC), and characterized by matrix-assisted laser desorption/ionization time-of-flight (MALDI-TOF) mass spectrometry.

### Plasmids, site-directed mutation, cell culture, and transient transfection

cDNAs encoding human K_V_1.1(homo sapiens), K_V_1.2(homo sapiens), K_V_1.3(homo sapiens), K_V_1.4(rattus norvegicus), K_V_1.5(homo sapiens), K_V_1.6(mus musculus), K_V_1.7(homo sapiens), K_V_2.1(rattus norvegicus), K_V_3.4(homo sapiens), K_V_4.1(rattus norvegicus), K_V_4.2(mus musculus), K_V_4.3(mus musculus), K_V_7.2(homo sapiens), Ca_V_1.2(rattus norvegicus), Ca_V_2.2(rattus norvegicus), Ca_V_3.1(mus musculus), Na_V_1.3(rattus norvegicus), Na_V_1.4(rattus norvegicus), Na_V_1.5(homo sapiens), Na_V_1.7(homo sapiens), and Na_V_1.8(mus musculus) channels were subcloned into pcDNA3.1 or pCMV-Blank mammalian expression vectors and maintained in our laboratory. Chimeric and mutant channels were generated by site-directed mutagenesis as described previously. All constructs were verified by Sanger sequencing.

HEK293T and ND7/23 cells were maintained in Dulbecco’s Modified Eagle Medium (DMEM) supplemented with 10% fetal bovine serum (FBS) and 1% penicillin–streptomycin (PS) (all from Gibco, Thermo Fisher Scientific) at 37 L in a humidified atmosphere containing 5% CO_2_. For electrophysiological recordings, wild-type or engineered K_V_, Na_V_, and Ca_V_ channels were transiently expressed in HEK293T or ND7/23 cells using Lipofectamine 2000 (Invitrogen, Thermo Fisher Scientific) according to the manufacturer’s instructions. Channel α-subunits, together with the appropriate auxiliary subunits when required, were co-transfected with pEGFP-N1 to identify transfected cells. Four to six hours after transfection, cells were replated onto poly-D-lysine-coated coverslips and cultured for an additional 24-36 h before electrophysiological analysis.

### Electrophysiology

Whole-cell patch-clamp recordings were performed using an EPC-10 amplifier controlled by PatchMaster software and analyzed with FitMaster (HEKA Elektronik). All experiments were conducted at room temperature. Whole-cell configuration was established using standard procedures. Only recordings with seal resistances >1 GΩ and series resistances <10 MΩ were included for analysis. Fast and slow capacitance transients were compensated electronically, and series resistance was compensated by 80% with a 100-μs lag to minimize voltage-clamp errors.

For K_V_ channel ionic-current recordings, the intracellular solution contained (mM): 140 KCl, 2.5 MgClL, 10 HEPES, and 11 EGTA (pH 7.4 with KOH), and the extracellular solution contained (mM): 145 NaCl, 2.8 KCl, 2 CaClL, 1 MgClL, 10 HEPES, and 10 D-glucose (pH 7.3 with NaOH). For K_V_1.3 gating-current recordings, the intracellular solution contained (mM): 140 NMDG, 50 HEPES, 50 HF, 5 EGTA, and 1 NMDG-Cl (pH 7.3 with methanesulfonic acid), whereas the extracellular solution contained (mM): 140 NMDG, 50 HEPES, 2 CaClL, 2 MgClL, 0.1 EDTA, and 0.01 CsCl (pH 7.3 with methanesulfonic acid). For Na_V_ current recordings, the intracellular solution contained (mM): 140 CsF, 10 NaCl, 10 HEPES, and 1 EGTA (pH 7.4 with CsOH), and the extracellular solution contained (mM): 140 NaCl, 5 KCl, 2 CaClL, 1 MgClL, 10 HEPES, and 10 D-glucose (pH 7.4 with NaOH). For Ca_V_ current recordings, the intracellular solution contained (mM): 120 CsMeSOL, 2 Mg-ATP, 10 HEPES, and 11 EGTA (pH 7.4 with CsOH), and the extracellular solution contained (mM): 138 NaCl, 20 TEA-Cl, 2.6 KCl, 2.6 CaClL, 1.2 MgClL, 5 HEPES, and 5 D-glucose (pH 7.4 with NaOH). Bath and pipette solutions were adjusted to an osmolarity of 300-330 mOsm and 290-310mOsm, repectively. Unless otherwise indicated, reagents were obtained from Sigma-Aldrich.

### Imiquimod-induced mouse psoriasis model

All animal procedures were conducted in accordance with the National Institutes of Health guidelines for the care and use of laboratory animals and approved by the Animal Care and Use Committee of the College of Medicine, Hunan Normal University. Male BALB/c mice (5–6 weeks old; Hunan SJA Laboratory Animal Co., Ltd., Changsha, China) were housed under controlled conditions (23 ± 3°C, 12-h light/dark cycle, lights on from 07:00 to 19:00) with free access to food and water. Psoriasis-like dermatitis was induced by daily topical application of 5% imiquimod (IMQ; Mingxin Pharmaceutical, Sichuan, China) to the dehaired left ear at 62.5 mg/cm² for 10 consecutive days (days 0–9). Mice were randomly assigned to receive saline, PAP-1 (25 mg/kg), or MPiN (25 mg/kg) by intraperitoneal injection once daily throughout the IMQ treatment period. Control mice received Vaseline and saline instead of IMQ and drug treatment. Ear thickness was measured daily, and disease severity was independently assessed by two blinded investigators using the Psoriasis Area and Severity Index (PASI) scoring system (0–4: 0, none; 1, mild; 2, moderate; 3, severe; 4, very severe). On day 9, mice were euthanized, and ear tissues were collected for hematoxylin and eosin (H&E) staining to determine epidermal thickness. Serum TNF-α levels were quantified by ELISA according to the manufacturer’s instructions (Proteintech Group, Wuhan, China).

### Modeling and simulations

Homology models of human K_V_1.3 in the C-type state (with and without MPiN) were constructed using MODELLER 1 based on the cryo-EM structure 9EEF, validated with ProCheck2, and energy-minimized. Each model was embedded in a 1-palmitoyl-2-oleoyl-sn-glycero-3-phosphocholine (POPC) bilayer at 300 K using the OPM3 database^54^ for membrane positioning, solvated with simple point charge (SPC) water, neutralized with counterions, and supplemented with 150 mM NaCl to mimic physiological conditions. The system was subjected to the default DESMOND relaxation protocol (gradual release of restraints from 10 K to 300 K under NVT and NPT ensembles) before production simulations^55^. All-atom conventional molecular dynamics (CMD) simulations (∼500 ns) were performed using the OPLS-2005 force field^56^ and smooth particle mesh Ewald for long-range electrostatics, with trajectories recorded every 200 ps. Protein-ligand interactions were analyzed with the DESMOND Simulation Interaction Diagram (SID) module. For enhanced sampling, metadynamics (MetaD) simulations (60 ns) were conducted after the same relaxation procedure, with Gaussian height, width, and deposition stride set to 0.03 kcal/mol, 0.05 Å, and 0.09 ps, respectively^57^. The free-energy surface (FES) was constructed from the accumulated bias potential, allowing the system to escape local minima and estimate relative free energies. All simulations were carried out on GPU-accelerated workstations.

### Data analysis

Data are presented as mean±SEM. *n* denotes the number of independent experimental replicates. Equal group sizes were used within each experiment. Data analysis was performed using Igor Pro 6.10A (WaveMetrics, Lake Oswego, OR, USA), OriginPro 2025 (OriginLab, Northampton, MA, USA), and GraphPad Prism 9 (GraphPad Software, La Jolla, CA, USA). Statistical significance was determined using paired or unpaired Student’s t-tests, or one-way ANOVA followed by Dunnett’s or Tukey’s multiple-comparison test, as appropriate. Differences were considered statistically significant at p < 0.05.

### Statistics and reproducibility

Statistical analyses were performed using two-tailed paired or unpaired t-tests, or one-way ANOVA followed by post-hoc Dunnett’s or Tukey’s tests, as appropriate. A p-value of less than 0.05 was considered statistically significant. Significance levels are indicated as follows: *p < 0.05, **p < 0.01, ***p < 0.001, and ****p < 0.0001. All repetitions were performed independently, the legend in the figure shows the specific repeated values (n values).

### Data availability

The data supporting this study are available from the corresponding authors on reasonable request. All data are included in the article and supplementary information files.

## Acknowledgments

This work was supported by grants from the National Natural Science Foundation of China (Grant Nos. 32571462 and U24A20366), the Youth Science Fund Project (Category A) of the Hunan Provincial Natural Science Foundation, China (Grant No. 2026JJ20032), the Scientific Research Program of Hunan Provincial Furong Laboratory, China (Grant No. 2025PT5040), and the National Key R&D Program of China (Grant No. 2025YFA1308900).

## Author contributions

C.T., M.R., and Y.Y. designed the experiments. G.L., X.Z., H.X., Z.W., Z.Z. (Zheyang Zhang), J.S., Z.Z. (Zixuan Zhang), Y.P., H.L., and X.H. performed the experiments. C.T., M.R.,Y.Y., and P.C. analyzed and interpreted the data. C.T., M.R., and Y.Y. supervised the study. C.T. and Y.Y. wrote the manuscript.

## Competing interests

The authors declare no competing interests.

## Additional information

The manuscript contains supplementary information files.

## Notes

### Competing Interest Statement

The authors have declared no competing interest.

