## Supplementary information for "An allosteric pocket in K_V_1.3 defines a distinct chemical space for immunomodulator design"

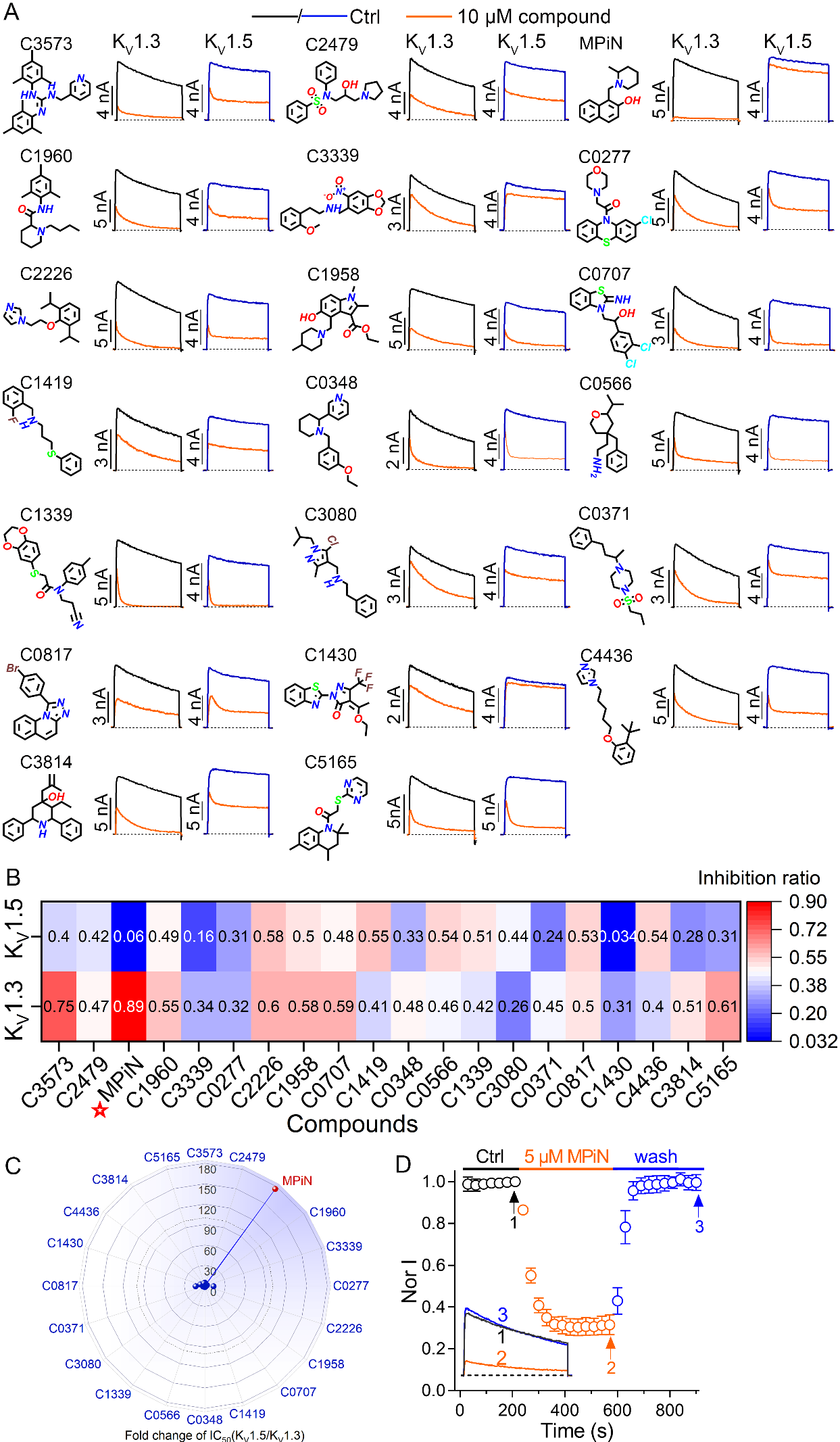

**Supplementary Figure 1.** (A) Typical K_V_1.3 and K_V_1.5 current traces before and after treatment with the indicated compounds, identified by screening K_V_1.3-active antagonists from a library containing 4,208 structurally diverse compounds (from Selleck Chemicals LLC, Catalog No. L3600) (n = 3-5). Currents were elicited using +20 mV/300 ms depolarizing pulses from a holding potential of -80 mV, with sweep intervals of 5 s. Compounds were applied by acute perfusion. (B) Heat map summarizing the compounds’ mean inhibition ratio on K_V_1.3 and K_V_1.5 currents, with the color scale (blue to red) representing low to high inhibition (n = 3-5). MPiN stood out for its potent inhibition of K_V_1.3 (0.89) with little effect on K_V_1.5 (0.06). (C) Fold increase in IC_50_ for inhibition of K_V_1.5 by the indicated compounds, normalized to the corresponding IC_50_ for K_V_1.3. IC_50_ values were calculated from the inhibition ratios in (B) using a simplified Hill equation (n = 3-5). (D) Time course of inhibition of K_V_1.3 currents by acute perfusion of 5 μM MPiN and recovery upon washout, with inhibition and recovery time constants of 24.8 ± 5.5 s and 35.7 ± 3.9 s, respectively (n = 7). Currents were normalized to the maximum current before MPiN administration (Ctrl). Inset shows typical currents at indicated time points.

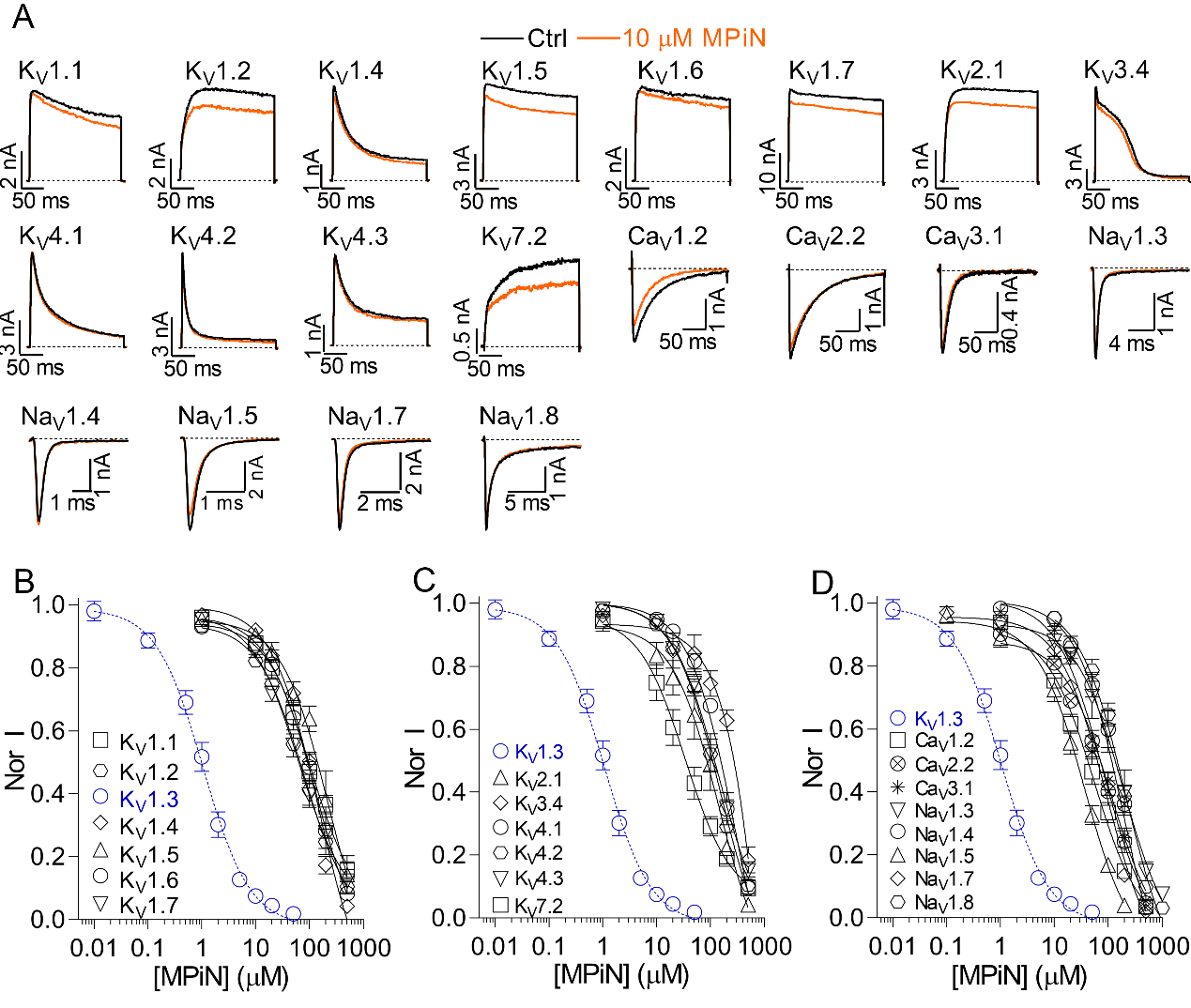

**Supplementary Figure 2.** MPiN selectively and potently inhibited the K_V_1.3 channel. (A) Representative current traces showing the effects of MPiN (10 μM) on various K_V_, Na_V_, and Ca_V_ channels as illustrated (n = 5-9). (B) - (D) Dose-response relationships of MPiN inhibiting K_V_1.3 and the off-target channels in (A) (n= 5-9). Currents with various doses of MPiN treatment were normalized to control current (Nor I). The IC_50_ values were summarized in **Supplementary Table 1**.

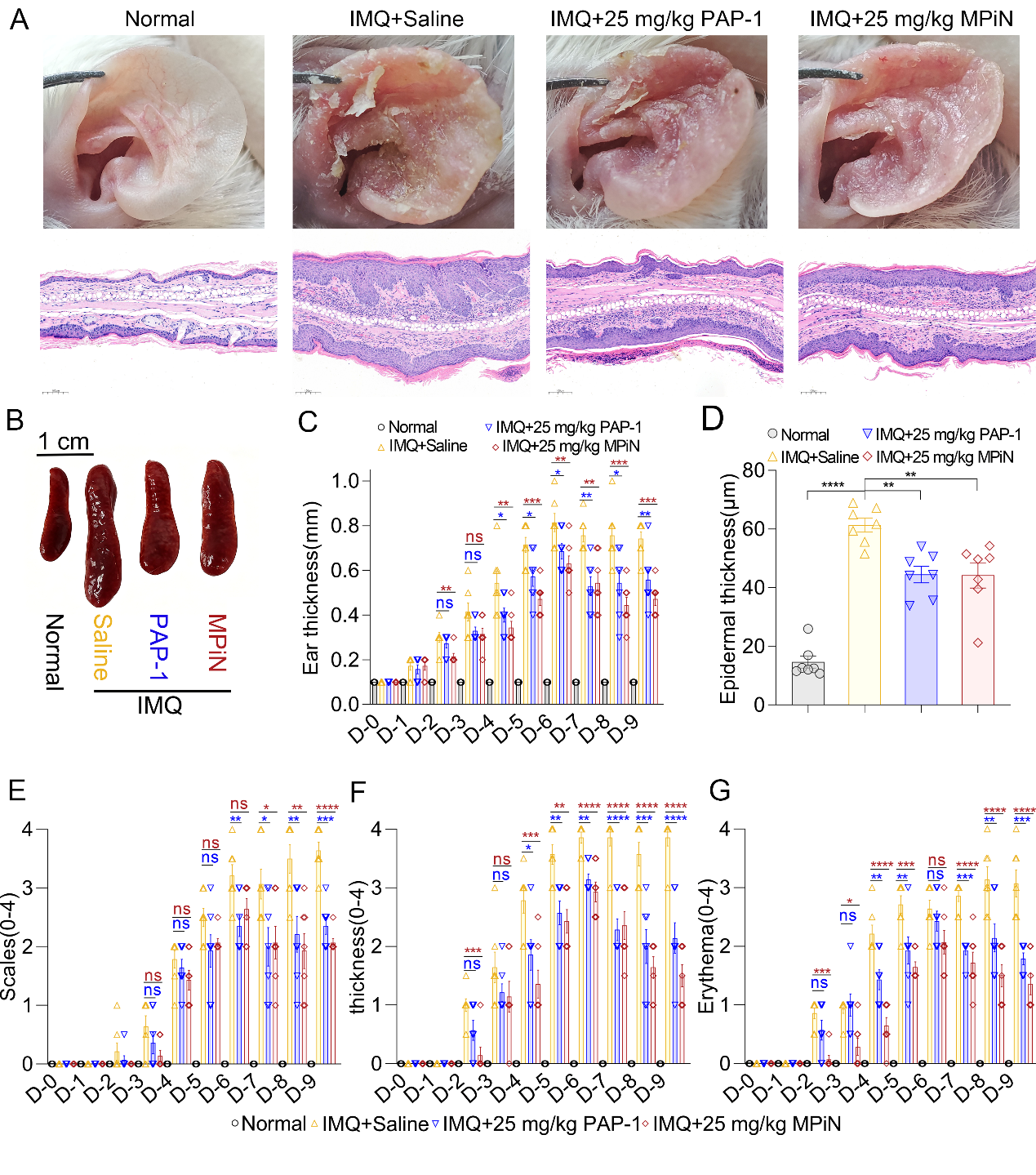

**Supplementary Figure 3.** MPiN alleviated IMQ-induced psoriasis in mice, with efficacy comparable to PAP-1. (A) Representative gross images of mouse ears (top) and H&E-stained ear sections (bottom) from day 9 (D9) mice (n= 7). (B) Representative spleen images for animals in each group (n= 7). (C) Time course of ear thickness during treatment (n= 7). (D) Quantification of epidermal thickness in ear sections from D9 mice (n= 7). (E-G) Psoriasis Area and Severity Index (PASI) scores. Scores for scaling (E), erythema (F), and ear thickening (G) were graded on a scale of 0–4 (n= 7).

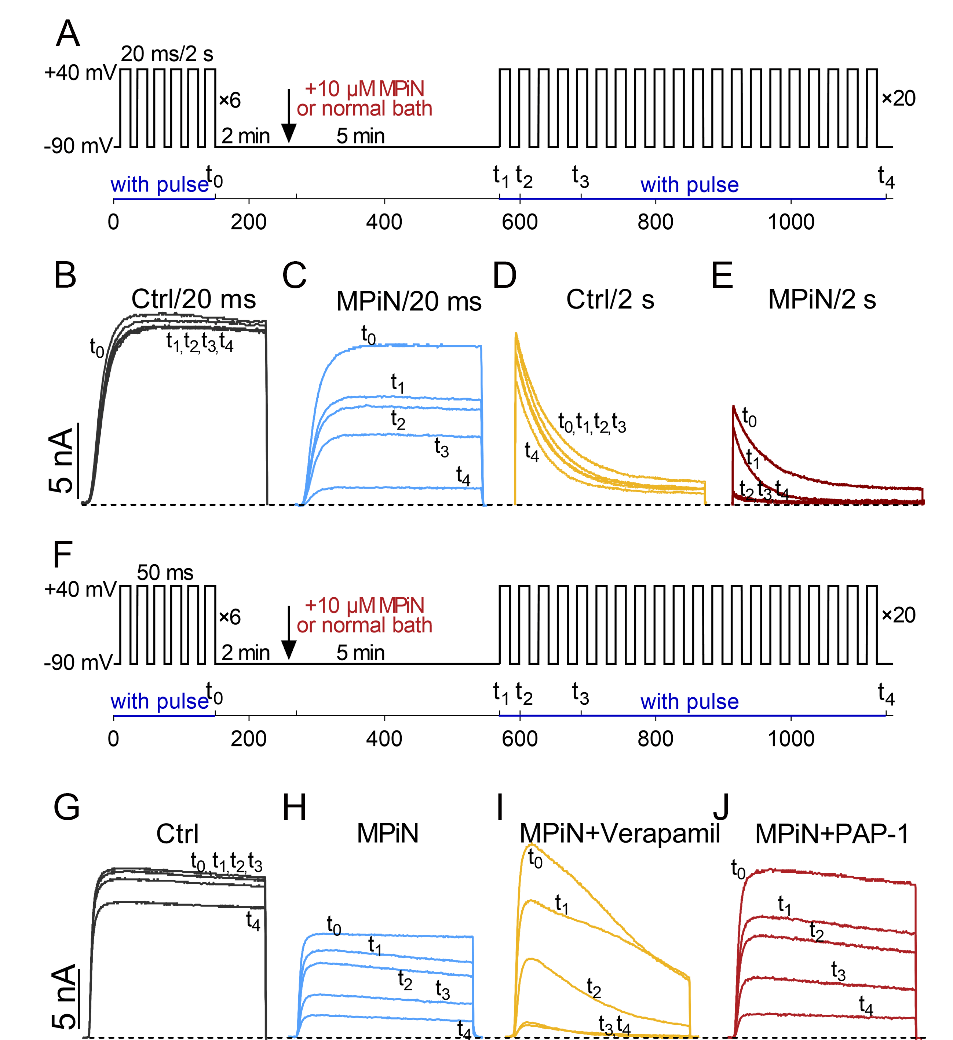

**Supplementary Figure 4.** (A) Voltage and drug perfusion protocols for analyzing the time course of K_V_1.3 current inhibition by MPiN. Channels were held at -90 mV and activated by two sets of depolarizing pulse trains (consisting of +40 mV depolarizing pulses of 20 ms or 2 s), separated by a -90 mV/7 min holding pulse that trapped channels in the resting state. The timeline represents a cumulative sweep interval of 30 s, with t_0_-t_4_ indicating different time points along the time course. 10 μM MPiN was applied by acute perfusion 2 min into the holding pulse, and incubated with resting state channels for an additional 5 min (red arrow) before depolarizing pulses were recovered. In the control group, normal bath solution was perfused instead. (B)-(E) Representative K_V_1.3 currents at the t_0_-t_4_ time points indicated in (A) (n = 8-11). (B, D) control group; (C, E) MPiN group. Currents were elicited by +40 mV pulses of 20 ms (B, C) or 2 s (D, E). (F) The same as in (A), with the exception of pulse duration of 50 ms. (G–J) Representative K_V_1.3 currents recorded at the t_0_–t_4_ time points indicated in (F) (n = 6-13). (G) Control; (H) MPiN; (I) verapamil + MPiN; and (J) PAP-1 + MPiN. Verapamil (100 μM) and PAP-1 (200 nM) were applied by preincubation from the beginning of the experiment.

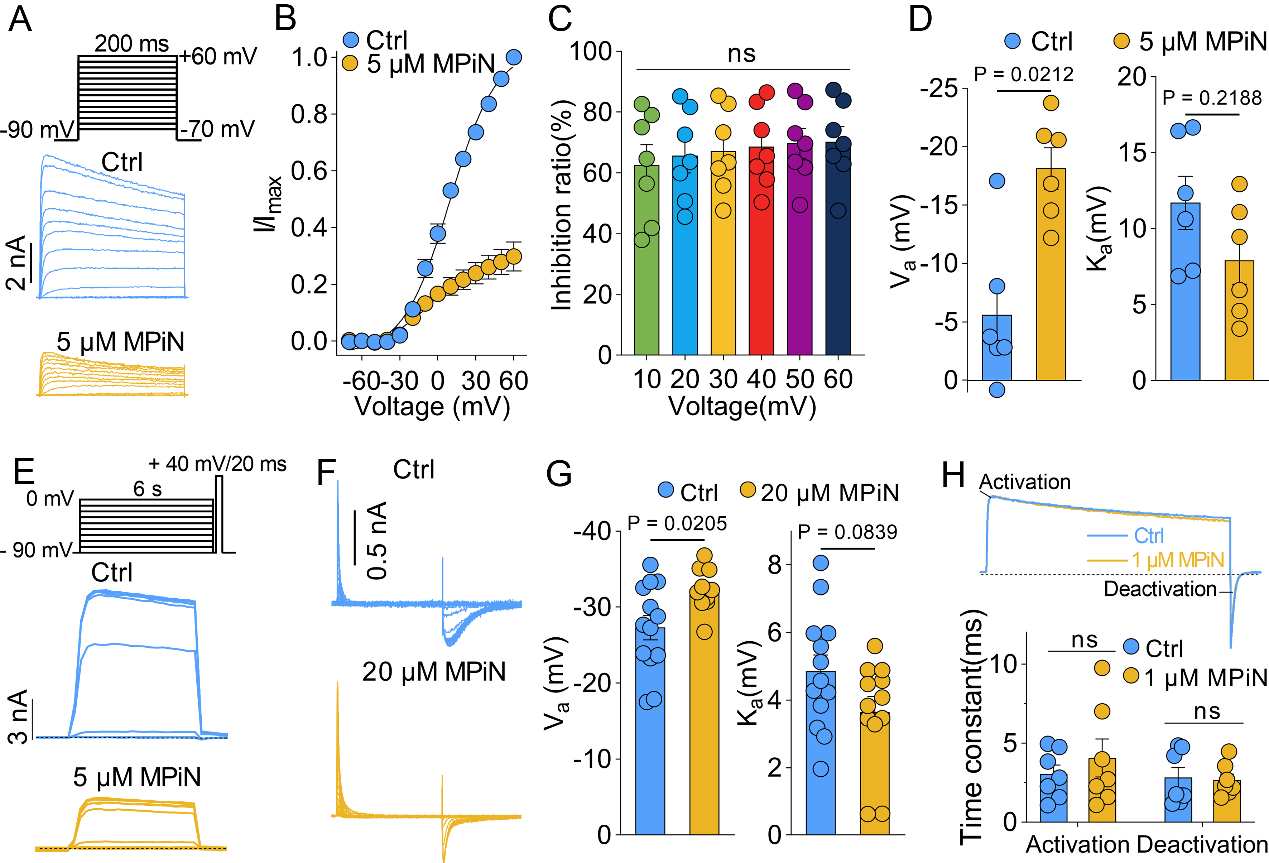

**Supplementary Figure 5.** (A) Representative K_V_1.3 currents before (blue) and after (yellow) 5 μM MPiN treatment (n = 6). Currents were evoked by depolarizing pulses from -70 mV to +60 mV (in 10 mV increment), from -90 mV holding, with sweep interval of 30 s. (B) Current-voltage (I-V) relationships from (A), with currents being normalized to the control current at +60 mV (n = 6). (C) Inhibition ratios across various depolarizing voltages in (B), showing voltage-independent inhibition by MPiN (n = 6). (D) Summary of half-activation voltage (V_a_, left panel) and slope factor (K_a_, right panel) for steady-state activation of K_V_1.3 channels before and after 5 μM MPiN treatment. Untreated: V_a_ = -6.9 ± 1.1 mV, K_a_ = 12.0 ± 1.0 mV; treated: V_a_ = -22.9 ± 1.0 mV, K_a_ = 6.4 ±0.9 mV. (E) Representative K_V_1.3 currents evoked using the classic two-pulses protocol shown, in the absence (blue) or presence (yellow) of preincubated MPiN (5 μM) (n = 6). (F) Typical K_V_1.3 gating currents elicited by depolarizations from -60 mV to +40 mV (in 10 mV increments), under conditions without or with preincubated MPiN (20 μM) (n = 12). (G) Summary of V_a_ (left panel) and K_a_ (right panel) for steady-state activation of K_V_1.3 gating currents in the absence and presence of 20 μM MPiN. Untreated: V_a_ = -29.2 ± 0.7 mV, K_a_ = 5.2 ± 0.5 mV; treated: V_a_ = -33.2 ± 0.4 mV, K_a_ = 4.3 ± 0.3 mV (n = 12). (H) Upper panel: representative K_V_1.3 currents before and after 1 μM MPiN treatment, each normalized to its own peak; Lower panel: activation and deactivation time constants calculated by fitting the current activation and deactivation processes, confirming no change in current kinetics. Untreated: τ_activation_ =3.0 ± 0.5 ms, τ_deactivation_ = 3.0 ± 0.6 ms; treated: τ_activation_ = 4.1 ± 1.1 ms , τ_deactivation_ = 2.8 ± 0.3 ms (n = 7). In (C), (D), (G), and (H), each dot represents a separate experimental cell. Statistical tests: (C) one-way ANOVA with post-hoc Tukey test; (D) and (H) paired t-test; (G) unpaired t-test.

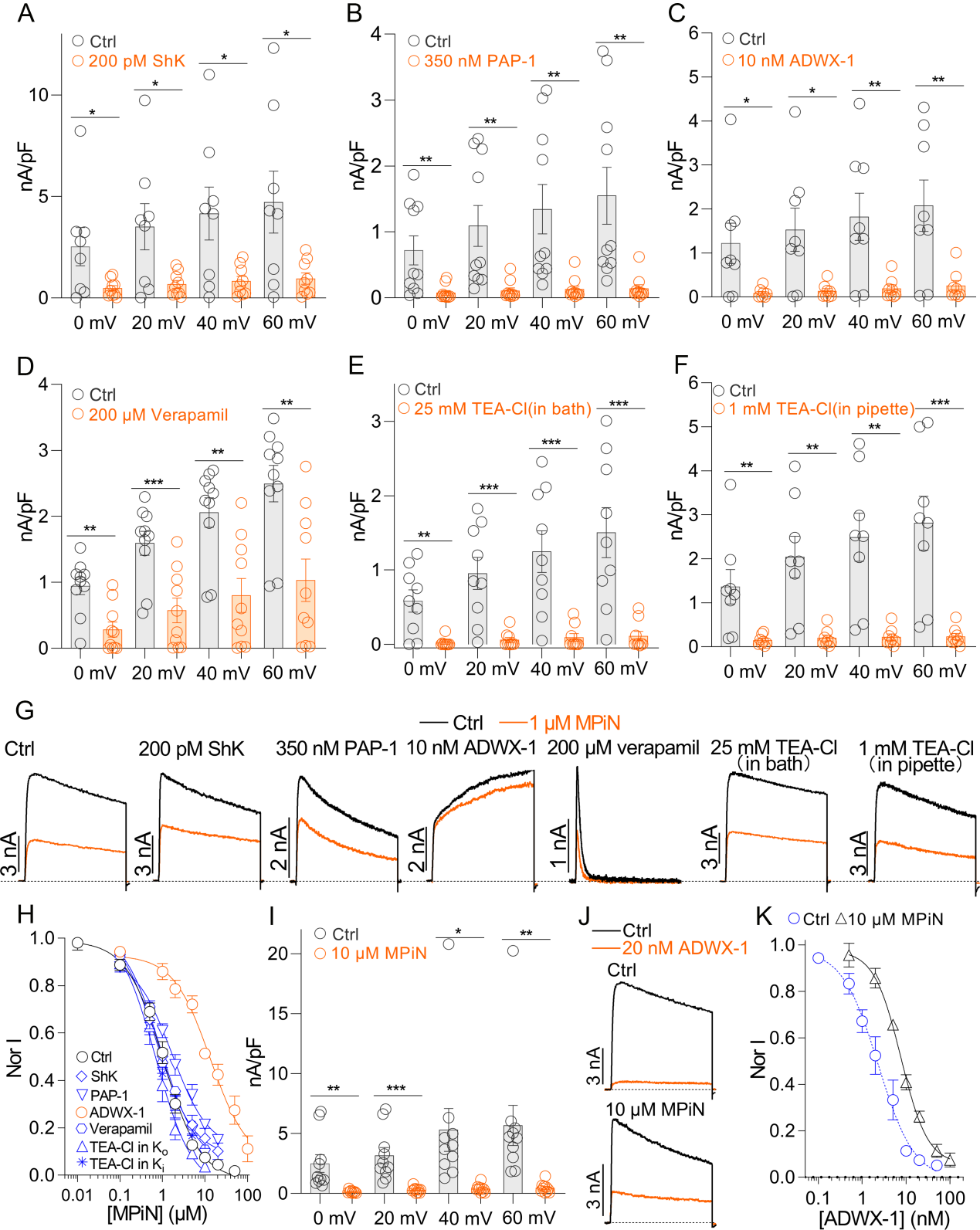

**Supplementary Figure 6.** (A-F) Current density of cells expressing K_V_1.3 channels across various depolarizing voltages, recorded without (Ctrl) or with bath preincubation of 200 pM ShK (A), 350 nM PAP-1 (B), 10 nM ADWX-1 (C), 200 μM verapamil (D), 25 mM TEA-Cl (E), or with 1 mM TEA-Cl in the pipette (F). Each dot represents a separate experimental cell, statistical differences were assessed by unpaired t-test (n = 8-10). (G) Representative current traces showing K_V_1.3 inhibition by 1 μM MPiN in the presence of the bath- or pipette-applied drugs shown in (A - F) (n = 8-10). (H) Dose-response relationships of MPiN inhibiting K_V_1.3 currents in the absence (Ctrl) or presence of preincubated drugs as indicated (n = 5-8). (I) Current density of K_V_1.3-expressing cells in the absence (Ctrl) or presence of 10 μM MPiN preincubation. Each dot represents an individual cell. Statistical significance was assessed by an unpaired t test (n = 10). (J - K) Representative current traces (J) and concentration-response curves (K) for ADWX-1 inhibition of K_V_1.3 channels in the absence (Ctrl) or presence of MPiN preincubation (n = 8). The IC_50_ values were summarized in **Supplementary Table 1**.

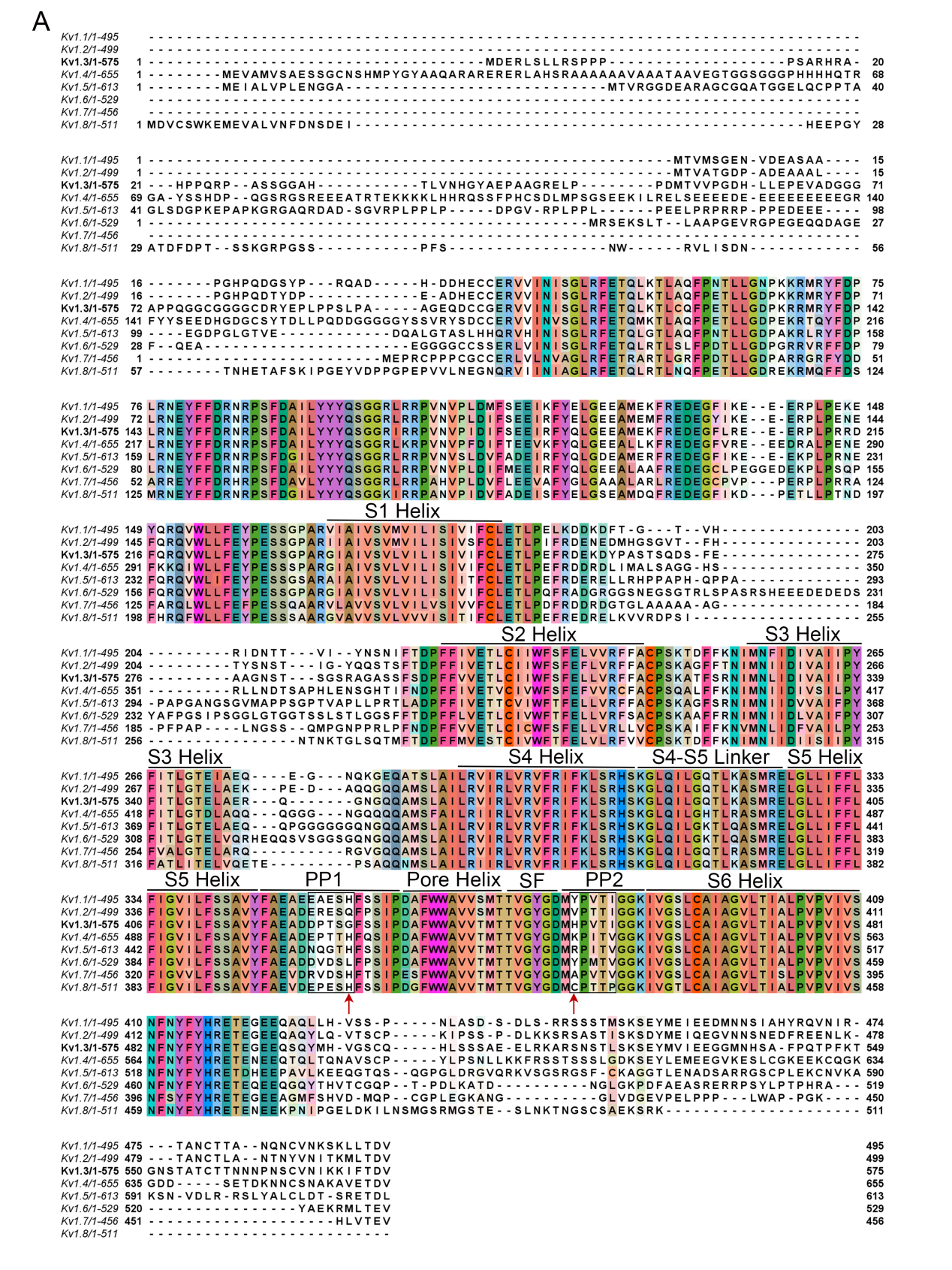

**Supplementary Figure 7.** Sequence alignment of K_V_1.x paralogs, with transmembrane helices S1-S6 and key linkers labeled: the extracellular S1-S2 and S3-S4 linkers, and the S5-S6 linker. The S5-S6 linker comprises the S5-pore helix loop (PP1 loop), the pore helix, the selectivity filter (SF), and the SF-S6 loop (PP2 loop).

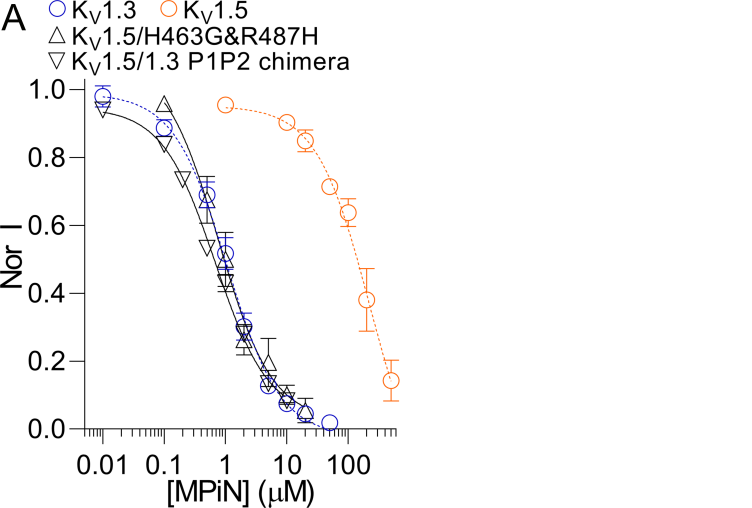

**Supplementary Figure 8.** Dose-response relationships of MPiN inhibiting the K_V_1.5/1.3 P1P2 chimera and K_V_1.5-H463G/R487H mutant channels, with K_V_1.3 and K_V_1.5 shown as dashed lines for reference (n = 6-7). The IC_50_ values were summarized in **Supplementary Table 1**.

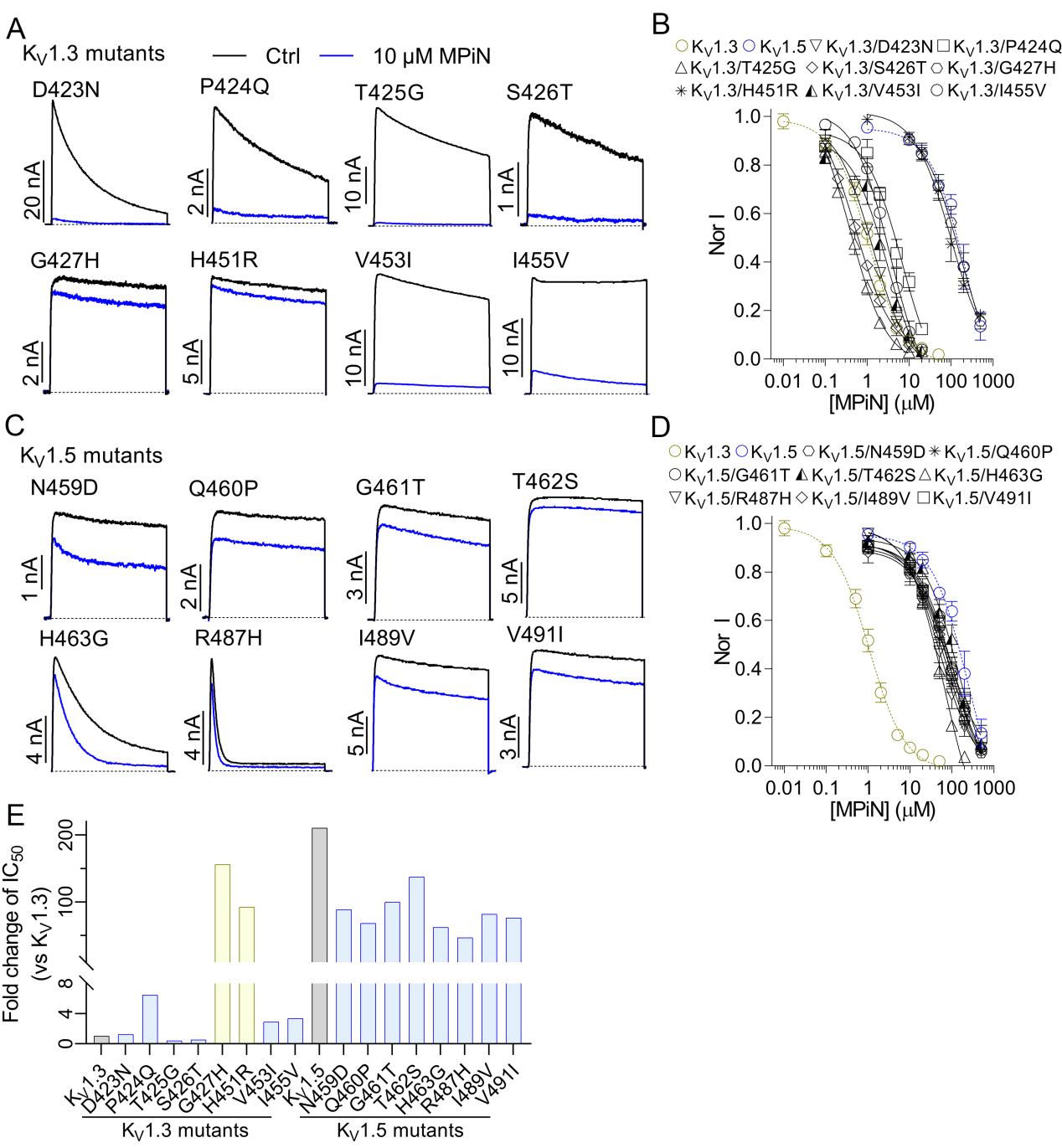

**Supplementary Figure 9.** (A) and (C) Representative current traces showing inhibition by 10 μM MPiN of (A) K_V_1.3- and (C) K_V_1.5-derived mutant channels, constructed using scanning mutagenesis to exchange variable residues in their S5-6 region (n = 5-9). (B) and (D) Dose-response relationships for MPiN inhibition of these (A) K_V_1.3- and (C) K_V_1.5-derived mutant channels; wild-type K_V_1.3 and K_V_1.5 channels were shown as dashed lines for reference (n =5-9). The IC_50_ values were summarized in **Supplementary Table 1**. (E) Fold change in IC_50_ for MPiN inhibition of these mutant channels relative to wild-type K_V_1.3 (n = 5-9).

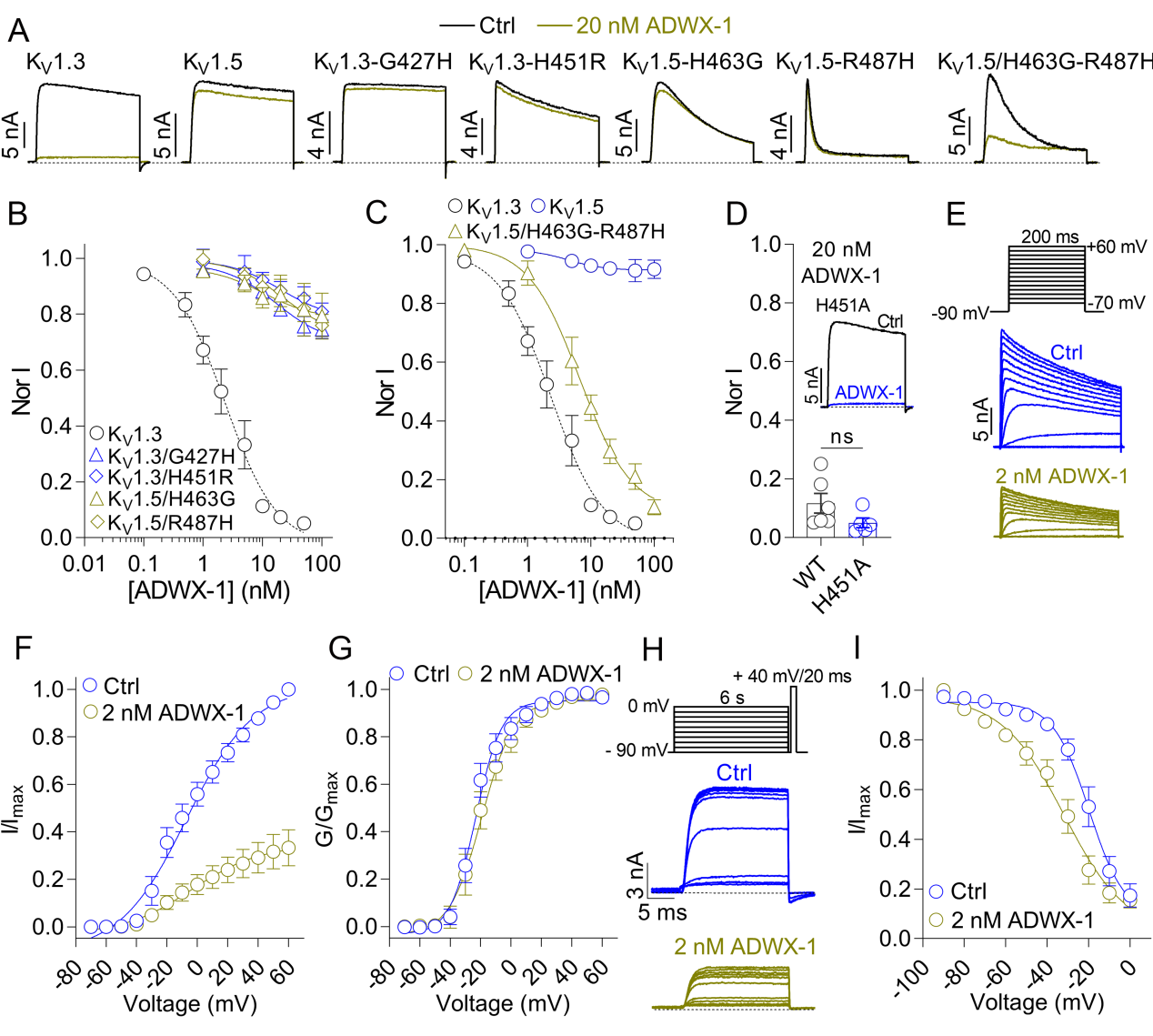

**Supplementary Figure 10.** (A-C) Representative current traces (A) and dose-response relationships (B, C) showing inhibition of K_V_1.3, K_V_1.5, K_V_1.3/G427H, K_V_1.3/H451R, K_V_1.5/H463G, K_V_1.5/R487H, and K_V_1.5/H463G/R487H channels by ADWX-1 (n =5-9). Substitution of G427 or H451 in K_V_1.3 with the corresponding K_V_1.5 residues (H463 and R487, respectively) drastically reduced ADWX-1 sensitivity. Conversely, ADWX-1 sensitivity was restored only by the double reverse mutation (H463G/R487H) in K_V_1.5. IC_50_ values were summarized in **Supplementary Table 1**. (D) Summary of normalized residual current ratio following treatment with 20 nM ADWX-1 in wild-type K_V_1.3 and K_V_1.3/H451A channels (n = 5-6). Inset depicts the typical current trace. (E) Representative K_V_1.3 currents recorded before (blue) and after (yellow) 2 nM ADWX-1 treatment (n = 10). Currents were elicited by depolarizing steps from -70 to +60 mV in 10-mV increments from a holding potential of -90 mV, with a sweep interval of 30 s. (F, G) Current-voltage (I - V; F) and conductance-voltage (G - V; G) relationships of K_V_1.3 channels under the conditions shown in (E). In (F), currents were normalized to the control current at +60 mV (I/I_Ctrl, +60 mV_). In (G), conductance values were normalized to the maximal conductance of each condition (G/G_max_). (H) Representative K_V_1.3 currents evoked using the two-pulse protocol shown, in the absence (blue) or presence (yellow) of 2 nM ADWX-1 preincubation (n = 10). (I) Steady-state fast inactivation curves of K_V_1.3 channels under the conditions shown in (H). The half-inactivation voltage (V_h_) and slope factor (K_h_) were -19.8 ± 2.5 mV and -8.6 ± 1.8 mV, and -31.8 ± 3.9 mV and -15.0 ± 3.6 mV, for ADWX-1 untreated and treated channels, respectively (n = 10).

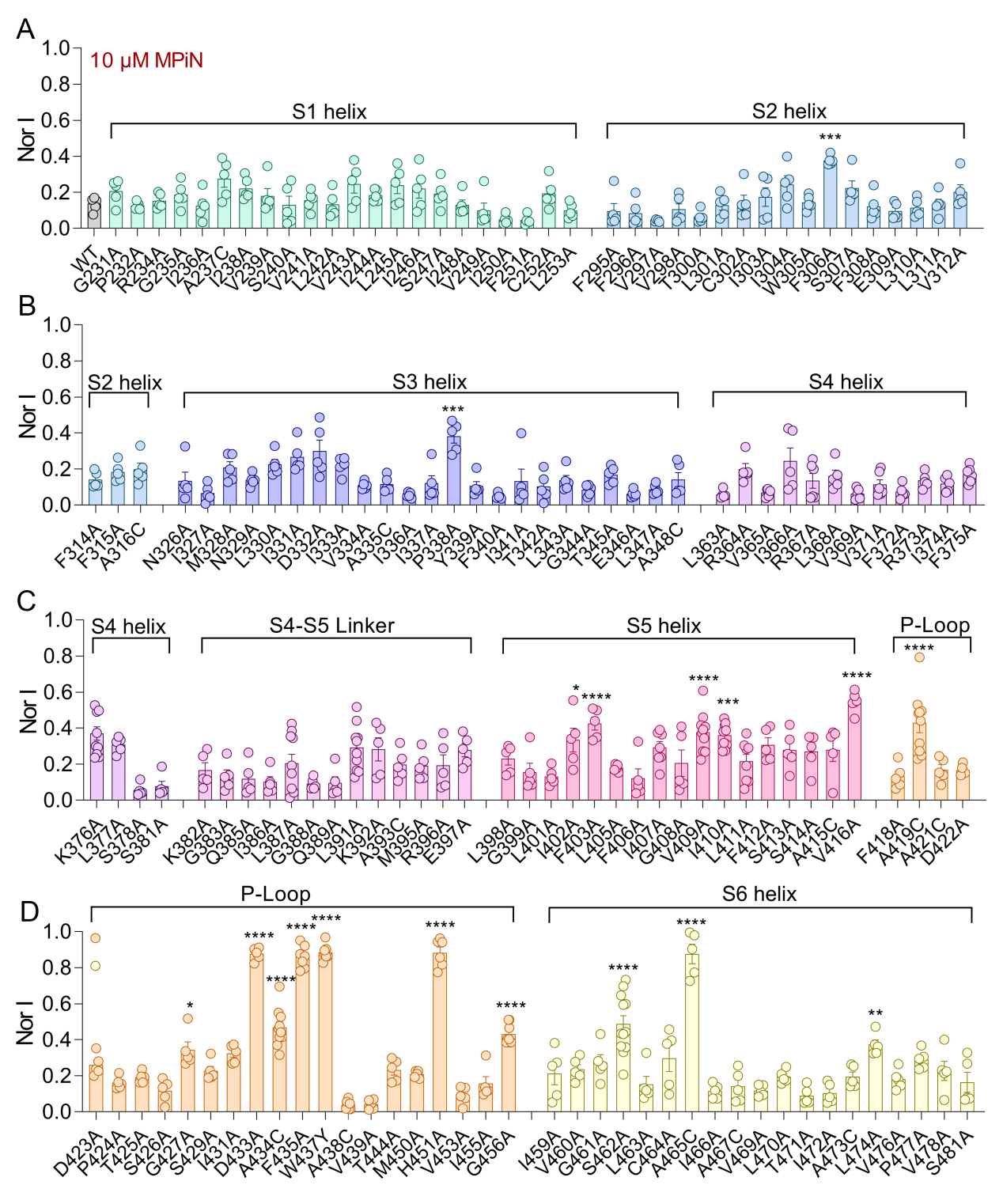

**Supplementary Figure 11.** Summary of normalized residual current ratios after 10 μM MPiN treatment for K_V_1.3 mutants generated by alanine-scanning mutagenesis (or cysteine substitution for native alanine residues) across the full-length K_V_1.3 sequence, excluding the intracellular loops and the S1-S2 and S3-S4 extracellular linkers (n = 5-10). Statistics: *p < 0.05; ***p < 0.001; ****p < 0.0001; one-way ANOVA with Dunnett's post-hoc test (wild-type K_V_1.3 as control).

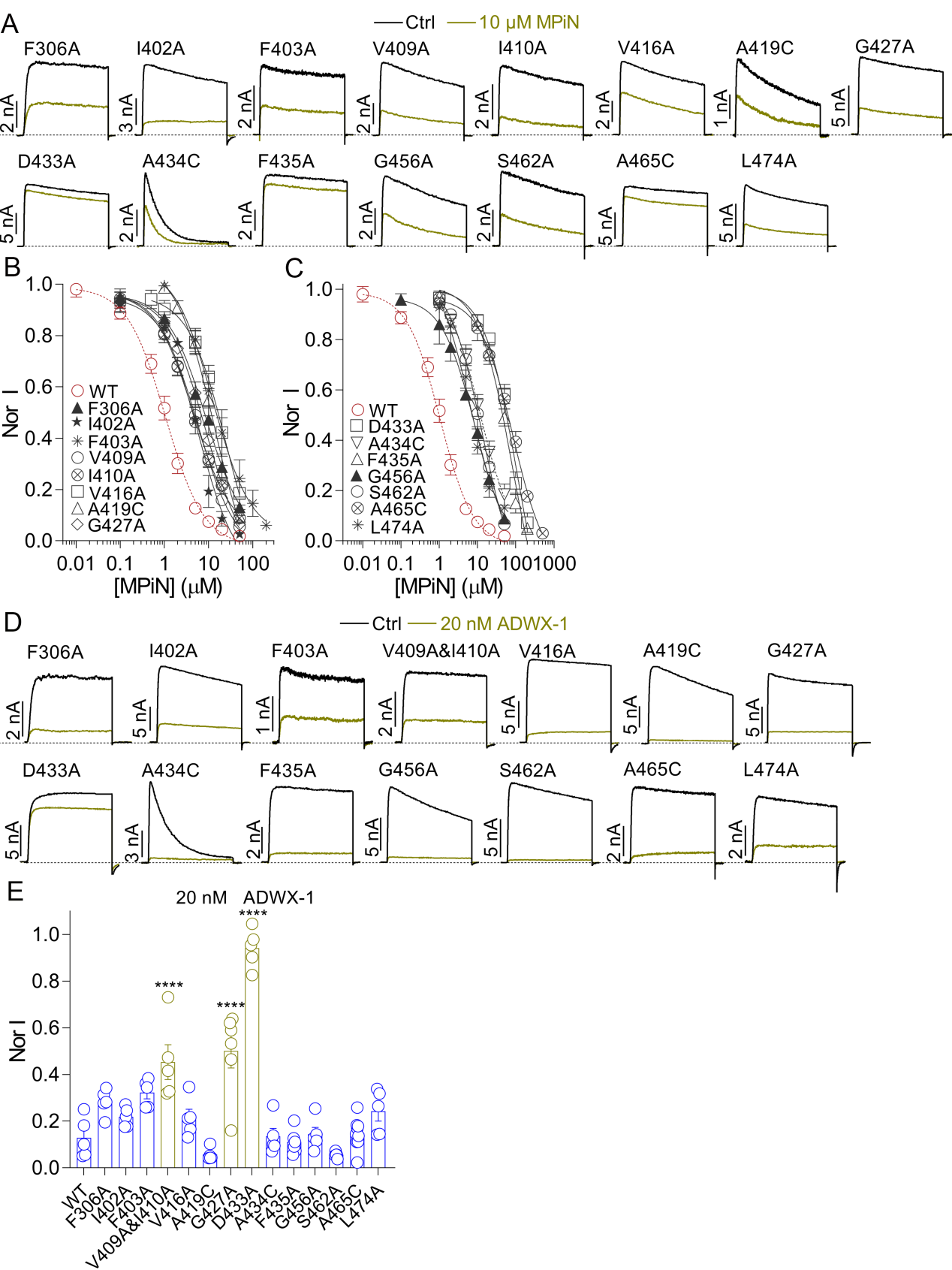

**Supplementary Figure 12.** (A-C) Representative current traces (A) and dose-response curves (B-C) showing MPiN inhibition of K_V_1.3 mutant channels identified by alanine/cysteine-scanning mutagenesis as critical for MPiN activity (Supplementary Figure 12; n = 5-7). Wild-type K_V_1.3 is shown for comparison. IC_50_ values are summarized in **Supplementary Table 1**. (D-E) Representative current traces (D) and summary of normalized residual current ratio (I_res_; E) showing inhibition by 20 nM ADWX-1 of these MPiN-critical mutant channels. Wild-type K_V_1.3 served as a reference. Each dot in (E) represents an individual cell (n = 5-8); significance was assessed by one-way ANOVA with Dunnett’s post hoc test. Note I_res_ values determined in (E) were used to calculate IC_50_s of ADWX-1 against K_V_1.3 and K_V_1.3 mutant channels based on the simplified Hill equation: IC_50_ = [X]/(1/I_res_ - 1), in which [X] was the ADWX-1 concentration. The calculated ADWX-1 IC_50_ for K_V_1.3 (2.9 ± 0.8 nM) was nearly identical to that obtained from the dose-response curve (2.3 ± 0.6 nM).

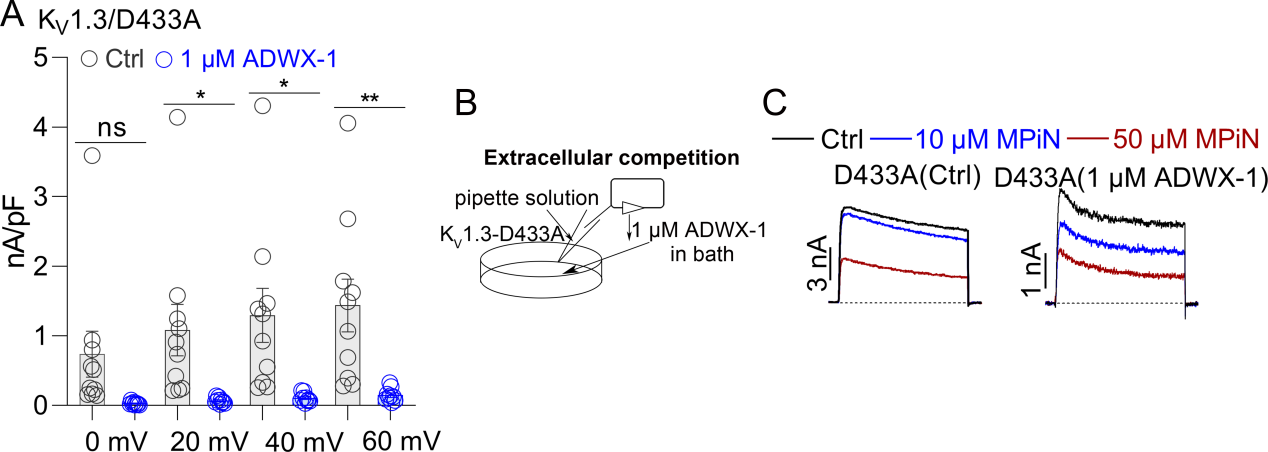

**Supplementary Figure 13.** (A) Current density-voltage relationships of cells expressing K_V_1.3/D433A mutant channels in the absence or presence of 1 μM ADWX-1 preincubation, showing robust channel inhibition. Each dot represents a separate experimental cell (n = 9-10). Significant difference was assessed by unpaired t-test. (B) Schematic of drug competition assay analyzing the influence of preincubated ADWX-1 on MPiN potency against the K_V_1.3/D433A mutant channel. (C) Representative current traces of MPiN inhibiting the K_V_1.3/D433A mutant channel in the absence or presence of bath-applied ADWX-1(n = 5-6).

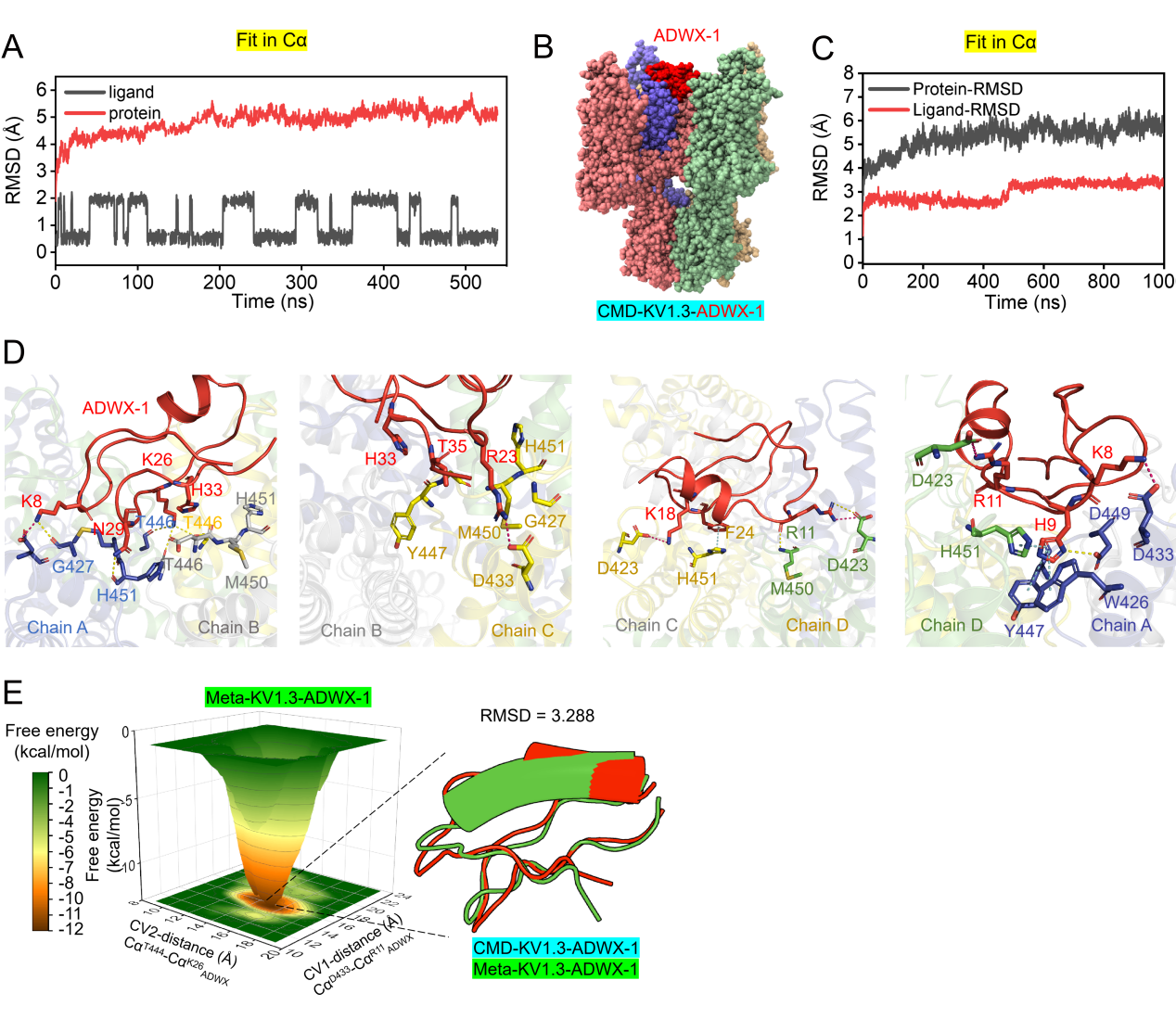

**Supplementary Figure 14.** **Binding of ADWX-1 to K_V_1.3 as observed by CMD and MetaD.** **(A)** RMSD plots of the ligand and protein during CMD simulation of the K_V_1.3–MPiN complex. **(B)** Front view of the three-dimensional complex of ADWX-1 bound to K_V_1.3. **(C)** RMSD plots of the ligand and protein during MetaD simulation of the ADWX-1–MPiN complex. **(D)** Detailed molecular structure of the protein–ligand interactions between ADWX-1 and K_V_1.3. **(E)** Conformation of the ADWX-1–Kv1.3 complex at the lowest free energy, as revealed by MetaD simulations.

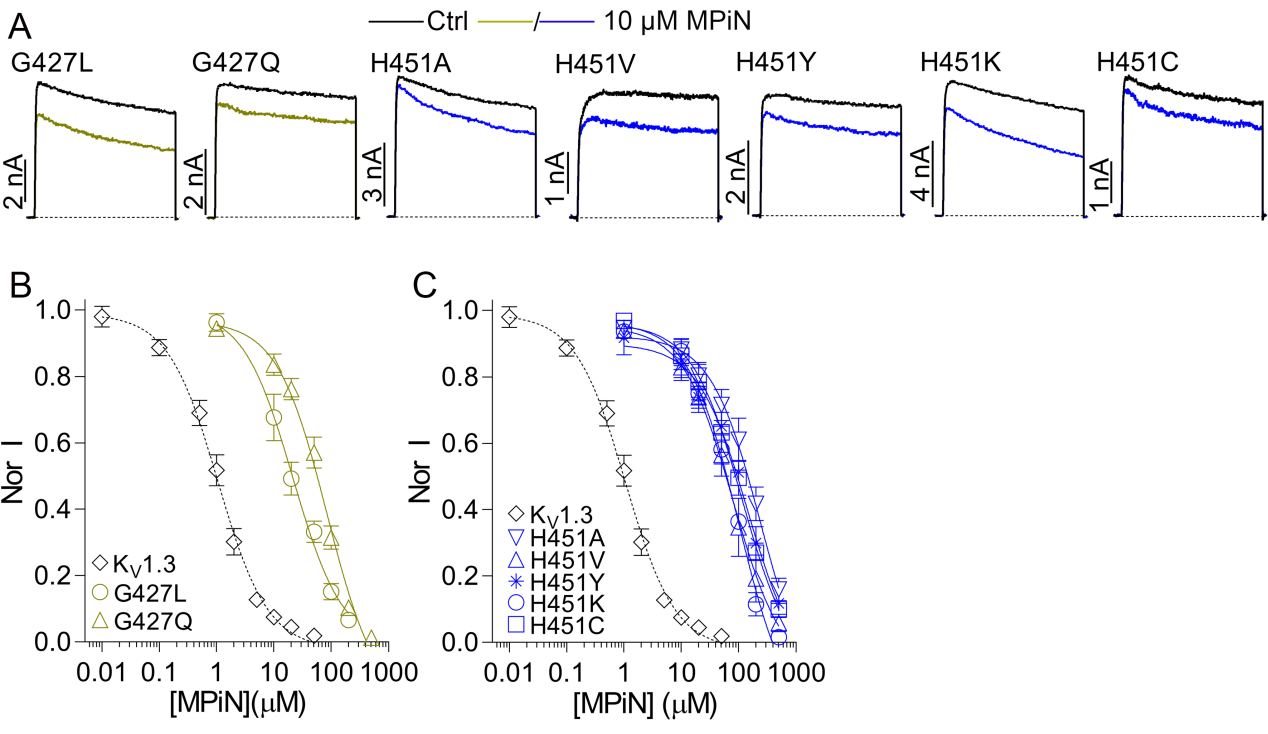

**Supplementary Figure 15.** (A) Representative current traces showing 10 μM MPiN inhibition of K_V_1.3 mutants, in which Gly427 (G427) or His451 (H451) was substituted with corresponding residues from K_V_1.1-1.2, K_V_1.4, or K_V_1.6-1.8 (n = 6-7). (B-C) Dose-response curves for MPiN inhibition of the (B) G427 and (C) H451 mutants, with wild-type K_V_1.3 and K_V_1.5 shown as dashed lines for reference (n = 6-7). The IC_50_ values were summarized in **Supplementary Table 1**.

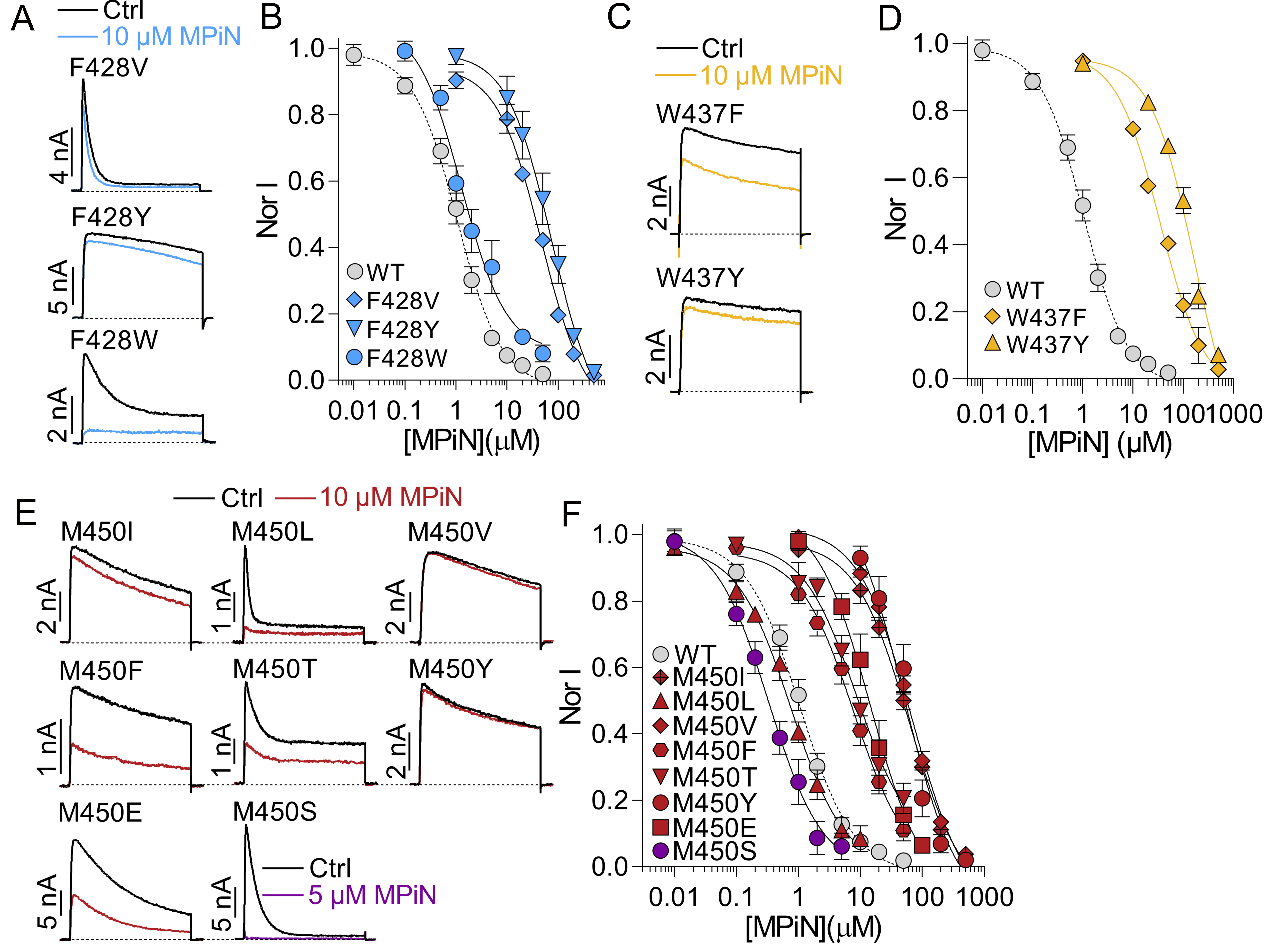

**Supplementary Figure 16.** Representative current traces (A, C, E) and concentration-response relationships (B, D, F) of MPiN inhibiting mutant channels generated by multiple-attribute mutations of the F428 (A-B), W437 (C-D), and M450 (E-F) residues in K_V_1.3 (n = 5-7). IC_50_ values were summarized in **Supplementary Table 1**.

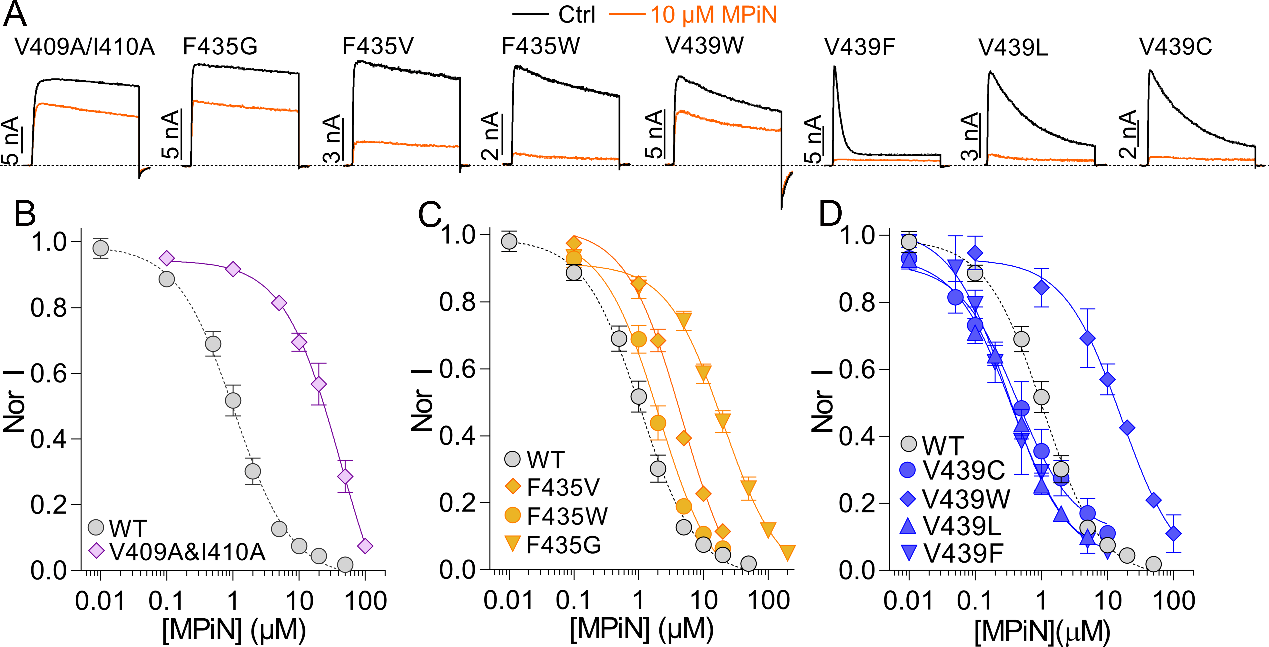

**Supplementary Figure 17.** Representative current traces (A) and concentration-response curves (B-D) showing MPiN inhibition of K_V_1.3 mutants bearing substitutions at residues along the putative allosteric propagation pathway (V409, I410, F435, and V439), with wild-type K_V_1.3 included for comparison (n = 6-10). IC_50_ values are summarized in **Supplementary Table 1**.

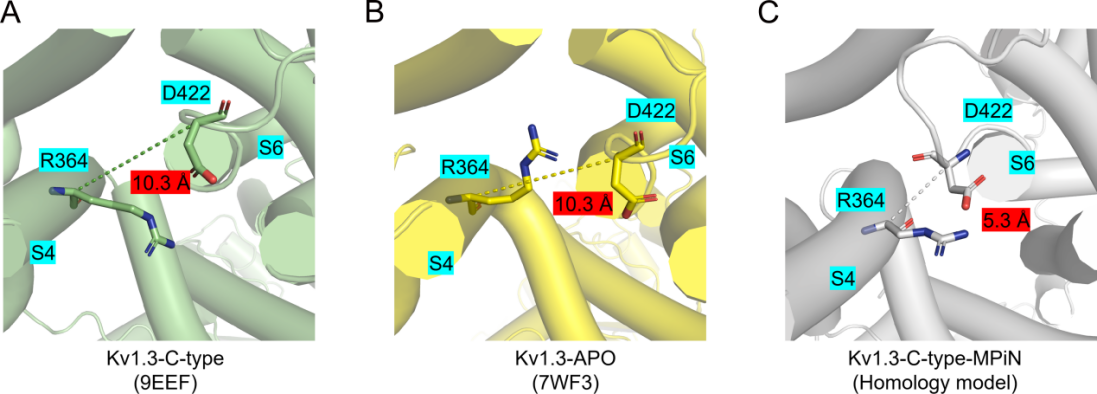

**Supplementary Figure 18.** **Interactions between the S6 and S4 regions of K_V_1.3 in distinct functional states. (A–C)** Shown are the interaction patterns and inter‑residue distances between R364 and D422 for K_V_1.3 in the C-Type inactivated state (A), the APO state (B), and the MPiN‑bound C-Type inactivated state (C), respectively.

**Supplementary Table 1**

Summary of IC_50_ values reported in this study

| **Figures** | **Channel** | **IC_50_ (μM)** | **n** | **Fold change of IC_50_**  **(*vs* K_V_1.3)** |
| --- | --- | --- | --- | --- |
| Fig.1B | K_V_1.3（-90 mV holding） | 1.05 ± 0.1 | 12 | 1 |
|  | K_V_1.3（-35 mV holding） | 0.6 ± 0.1 | 7 | 0.55 |
| Fig.1C/SFig.2 | K_V_1.1 | 101.2 ± 16.3 | 9 | 96.1 |
|  | K_V_1.2 | 79.5 ± 13.5 | 6 | 75.5 |
|  | K_V_1.4 | 150.6 ± 23.2 | 6 | 142.9 |
|  | K_V_1.5 | 225.1 ± 64.6 | 6 | 213.7 |
|  | K_V_1.6 | 135.0 ± 23.6 | 5 | 128.2 |
|  | K_V_1.7 | 63.4 ± 12.0 | 6 | 60.2 |
|  | K_V_2.1 | 144.0 ± 46.6 | 8 | 136.7 |
|  | K_V_3.4 | 336.6 ± 6.8^a^ | 5 | 319.7 |
|  | K_V_4.1 | 306.2 ± 48.4 | 6 | 290.7 |
|  | K_V_4.2 | 151.4 ± 17.6 | 6 | 143.8 |
|  | K_V_4.3 | 142.2 ± 28.7 | 6 | 135 |
|  | K_V_7.2 | 33.9 ± 7.4 | 5 | 32.1 |
|  | Na_V_1.3 | 186.9 ± 48.1 | 5 | 177.4 |
|  | Na_V_1.4 | 199.2 ± 39.5 | 6 | 189.2 |
|  | Na_V_1.5 | 36.3 ± 5.2 | 7 | 34.4 |
|  | Na_V_1.7 | 91.9 ± 12.6 | 6 | 87.2 |
|  | Na_V_1.8 | 210.9 ± 34.4 | 7 | 199.3 |
|  | Ca_V_1.2 | 54.3 ± 8.8 | 5 | 51.5 |
|  | Ca_V_2.2 | 110.5 ± 21.5 | 6 | 104.9 |
|  | Ca_V_3.1 | 81.8 ± 11.1 | 5 | 77.7 |
| Fig.3C/SFig.6H | K_V_1.3(with 200 pM ShK in bath) | 0.7 ± 0.2 | 7 | 1 |
|  | K_V_1.3(with 350 nM PAP-1 in bath) | 1.7 ± 0.4 | 6 | 0.7 |
|  | K_V_1.3(with 10 nM ADWX-1 in bath) | 13.3 ± 3.1 | 8 | 1.6 |
|  | K_V_1.3(with 200 μM verapamil in bath) | 0.9 ± 0.1 | 5 | 12.7 |
|  | K_V_1.3(with 25 mM TEA-Cl in bath) | 0.7 ± 0.2 | 6 | 0.8 |
|  | K_V_1.3(with 1 mM TEA-Cl in pipette) | 0.7 ± 0.1 | 6 | 0.7 |
| Fig.3D/SFig.6K | K_V_1.3 | 0.0023 ± 0.0006 | 8 | 1.0 |
|  | K_V_1.3(with 10 μM MPiN in bath) | 0.0076 ± 0.0009 | 8 | 3.3 |
| Fig.3F/SFig.8 | K_V_1.5/1.3 P1P2 chimera | 0.7 ± 0.1 | 7 | 0.6 |
|  | K_V_1.5-H463G/R487H | 0.8 ± 0.2 | 6 | 0.7 |
| Fig.3F/SFig.10C | K_V_1.5-H463G/R487H | 0.0066 ± 0.0014 | 9 | 2.9 |
|  | K_V_1.5 | >0.1 | 8 | >100 |
| Fig.4A  (upper panel)/SFig.12B-C | K_V_1.3/F306A | 8.7 ± 2.2 | 6 | 8.2 |
|  | K_V_1.3/I402A | 5.9 ± 1.3 | 5 | 5.6 |
|  | K_V_1.3/F403A | 12.5 ± 3.0 | 7 | 11.8 |
|  | K_V_1.3/V409A | 5.8 ± 0.9 | 5 | 5.5 |
|  | K_V_1.3/I410A | 4.5 ± 0.7 | 5 | 4.3 |
|  | K_V_1.3/V416A | 25.7 ± 9.5 | 7 | 24.4 |
|  | K_V_1.3/A419C | 13.7 ± 3.3 | 6 | 13.0 |
|  | K_V_1.3/G427A | 8.6 ± 0.9 | 5 | 8.1 |
|  | K_V_1.3/D433A | 111.7 ± 30.9 | 6 | 106.1 |
|  | K_V_1.3/A434C | 11.9 ± 3.0 | 6 | 11.3 |
|  | K_V_1.3/F435A | 81.1 ± 19.0 | 7 | 77.0 |
|  | K_V_1.3/W437Y | 157.5 ± 29.2 | 7 | 149.6 |
|  | K_V_1.3/G456A | 8.8 ± 1.8 | 5 | 8.3 |
|  | K_V_1.3/S462A | 13.7 ± 3.4 | 7 | 13.0 |
|  | K_V_1.3/A465C | 84.2 ± 16.0 | 5 | 80.0 |
|  | K_V_1.3/L474A | 6.3 ± 1.5 | 6 | 6.0 |
| Fig.4A  ( lower panel)/SFig.12E | K_V_1.3 | 0.0029 ± 0.0008^a^ | 5 | 1.0 |
|  | K_V_1.3/F306A | 0.0079 ± 0.0005^a^ | 5 | 2.7 |
|  | K_V_1.3/I402A | 0.0056 ± 0.0004^a^ | 5 | 1.9 |
|  | K_V_1.3/F403A | 0.0095 ± 0.0005^a^ | 5 | 3.2 |
|  | K_V_1.3-V409A/I410A | 0.0166 ± 0.0016^a^ | 5 | 5.6 |
|  | K_V_1.3/V416A | 0.0055 ± 0.0008^a^ | 5 | 1.9 |
|  | K_V_1.3/A419C | 0.0012 ± 0.0002^a^ | 5 | 0.4 |
|  | K_V_1.3/G427A | 0.0201 ± 0.0016^a^ | 6 | 6.8 |
|  | K_V_1.3/D433A | 0.3211 ± 0.0008^a^ | 5 | 109.0 |
|  | K_V_1.3/A434C | 0.0031 ± 0.0007^a^ | 5 | 1.1 |
|  | K_V_1.3/F435A | 0.0025 ± 0.0004^a^ | 6 | 0.8 |
|  | K_V_1.3/G456A | 0.0033 ± 0.0006^a^ | 5 | 1.1 |
|  | K_V_1.3/S462A | 0.0012 ± 0.0001^a^ | 5 | 0.4 |
|  | K_V_1.3/A465C | 0.0034 ± 0.0004^a^ | 8 | 1.2 |
|  | K_V_1.3/L474A | 0.0064 ± 0.0009^a^ | 5 | 2.2 |
| Fig.4C | K_V_1.3/D433A | 41.8 ± 4.4 | 5 | 39.7 |
|  | K_V_1.3/D433A(with 1 μM ADWX-1 in bath) | 40.4 ± 9.1 | 6 | 38.4 |
| Fig.4H/SFig.16B,D,F | K_V_1.3/F428Y | 74.6 ± 24.2 | 7 | 70.8 |
|  | K_V_1.3/F428V | 46.7 ± 5.9 | 5 | 44.3 |
|  | K_V_1.3/F428W | 1.3 ± 0.4 | 5 | 1.2 |
|  | K_V_1.3/W437Y | 166.1 ± 28.1 | 7 | 157.7 |
|  | K_V_1.3/W437F | 34.6 ± 4.1 | 6 | 32.8 |
|  | K_V_1.3/M450S | 0.3 ± 0.1 | 6 | 0.3 |
|  | K_V_1.3/M450Y | 50.8 ± 14.9 | 6 | 48.2 |
|  | K_V_1.3/M450T | 8.8 ± 2.4 | 7 | 8.3 |
|  | K_V_1.3/M450E | 14.7 ± 4.2 | 6 | 14.0 |
|  | K_V_1.3/M450F | 7.8 ± 1.7 | 6 | 7.4 |
|  | K_V_1.3/M450L | 0.8 ± 0.1 | 7 | 0.7 |
|  | K_V_1.3/M450V | 69.5 ± 6.6 | 6 | 65.9 |
|  | K_V_1.3/M450I | 61.6 ± 8.1 | 6 | 58.4 |
| Fig.5J/SFig.15B-C | K_V_1.3/G427L | 21.3 ± 6.1 | 6 | 20.3 |
|  | K_V_1.3/G427Q | 84.3 ± 14.1 | 7 | 80.1 |
|  | K_V_1.3/H451A | 240.1 ± 80.3 | 6 | 228.0 |
|  | K_V_1.3/H451V | 79.1 ± 20.1 | 6 | 75.1 |
|  | K_V_1.3/H451Y | 144.3 ± 34.9 | 6 | 137.0 |
|  | K_V_1.3/H451K | 100.4 ± 20.4 | 6 | 95.3 |
|  | K_V_1.3/H451C | 113.7 ± 26.3 | 6 | 108.0 |
| Fig.6E/SFig.17B-D | K_V_1.3-V409A/I410A | 43.1 ± 9.1 | 6 | 40.9 |
|  | K_V_1.3/F435W | 2.1 ± 0.4 | 6 | 2.0 |
|  | K_V_1.3/F435V | 4.8 ± 0.7 | 7 | 4.5 |
|  | K_V_1.3/F435G | 20.1 ± 2.7 | 7 | 19.1 |
|  | K_V_1.3/V439L | 0.4 ± 0.1 | 6 | 0.3 |
|  | K_V_1.3/V439W | 16.2 ± 4.9 | 6 | 15.3 |
|  | K_V_1.3/V439C | 0.4 ± 0.1 | 10 | 0.4 |
|  | K_V_1.3/V439F | 0.3 ± 0.1 | 8 | 0.3 |
| SFig.9B | K_V_1.3/D423N | 1.3 ± 0.2 | 7 | 1.3 |
|  | K_V_1.3/T425G | 0.4 ± 0.1 | 6 | 0.4 |
|  | K_V_1.3/S426T | 0.5 ± 0.1 | 5 | 0.5 |
|  | K_V_1.3/G427H | 166.9 ± 57.1 | 6 | 158.4 |
|  | K_V_1.3/H451R | 99.2 ± 30.8 | 9 | 94.2 |
|  | K_V_1.3/V453I | 3.1 ± 1.0 | 7 | 2.9 |
|  | K_V_1.3/P424Q | 6.9 ± 2.8 | 6 | 6.6 |
|  | K_V_1.3/I455V | 3.6 ± 0.9 | 5 | 3.4 |
| SFig.9D | K_V_1.5/H463G | 66.9 ± 16.3 | 8 | 63.6 |
|  | K_V_1.5/R487H | 50.4 ± 13.5 | 7 | 47.8 |
|  | K_V_1.5/I489V | 88.0 ± 15.9 | 6 | 83.6 |
|  | K_V_1.5/N459D | 95.0 ± 24.2 | 6 | 90.2 |
|  | K_V_1.5/T462S | 147.0 ± 37.3 | 6 | 139.6 |
|  | K_V_1.5/Q460P | 73.4 ± 11.5 | 7 | 69.6 |
|  | K_V_1.5/G461T | 107.0 ± 23.2 | 7 | 101.6 |
|  | K_V_1.5/V491I | 82.0 ± 17.1 | 6 | 77.8 |

^a^Estimated from a simplified Hill equation; all other IC_50_ values were determined by fitting the concentration-response curve.
